# Rapid and accurate multi-phenotype imputation for millions of individuals

**DOI:** 10.1101/2023.06.25.546422

**Authors:** Lin-Lin Gu, Hong-Shan Wu, Tian-Yi Liu, Yong-Jie Zhang, Jing-Cheng He, Xiao-Lei Liu, Zhi-Yong Wang, Guo-Bo Chen, Dan Jiang, Ming Fang

## Abstract

Deep phenotyping can enhance the power of genetic analysis, including genome-wide association studies (GWAS), but the occurrence of missing phenotypes compromises the potential of such resources. Although many phenotypic imputation methods have been developed, the accurate imputation of millions of individuals remains extremely challenging. In the present study, we developed a novel multi-<u>p</u>henotype imputation method based on mi<u>x</u>ed f<u>a</u>st ra<u>n</u>dom forest (PIXANT) by leveraging efficient machine learning (ML)-based algorithms. We demonstrate that PIXANT runtime is faster and computer memory usage is less than that of other state-of-the-art methods when applied to the UK Biobank (UKB) data, suggesting that PIXANT is scalable to cohorts with millions of individuals. Our simulations with hundreds of individuals showed that PIXANT accuracy was superior to or comparable to the accuracy of the most advanced methods available. PIXANT was used to impute 425 phenotypes for the UKB data of 277,301 unrelated White British citizens. When GWAS was subsequently performed on the imputed phenotypes, 18.4% more GWAS loci were identified than before imputation (8,710 *vs* 7,355). The increased statistical power of GWAS identified novel positional candidate genes affecting heart rate, such as *RNF220, SCN10A*, and *RGS6*, suggesting that the use of imputed phenotype data from a large cohort may lead to the discovery of novel genes for complex traits.

## Introduction

Large-scale data analysis, such as genome-wide association studies (GWAS), can significantly improve genetic research by increasing statistical power^1–3^. In recent years, we have witnessed tremendous growth in data available for GWAS. For example, the UK Biobank (UKB) database has collected nearly half a million genotyped individuals with enriched phenotypes^4^, and has played a crucial role in discovering novel genotype-phenotype associations in recent years^5–8^. The decreasing costs of high-throughput genotyping are expected to lead to the development of more expansive biobank datasets enriched with detailed phenotypic information. These large-scale datasets will, in turn, harness the scope and depth of genetic analyses. However, the move to high-dimensional phenotypes will inevitably lead to a higher missing rate. For example, the missing rate ranges from 0.11% to 98.35% in UKB, a loss that decreases the discovery rate in the downstream analyses^9^. If missing phenotypes can be imputed accurately, more complete datasets can be utilized for GWAS.

Traits often have a certain degree of correlation, which provides a scaffold for multiple-phenotype imputation. MICE is a commonly used multi-phenotype imputation method that uses chained equations to handle missing data^10^. The MICE method is popular with statisticians because of its efficient computation of large data. Compared with MICE, missForest has been developed and offers better performance when applied to continuous or binary traits^11^ but is computationally much slower^12,13^. Alternative variational Bayesian framework-based methods such as PHENIX have been proposed that better impute missing phenotypes^9^. PHENIX is based on a multivariate normal distribution containing a matrix of sample kinship and is more accurate than MICE and missForest; however, PHENIX computation is both time consuming and memory intensive^9^. It is estimated that PHENIX analysis of the UKB dataset would require several months of runtime and over 1T of computer memory. With continuous dataset expansion, the computational time and computer memory required will also increase, ultimately draining the computing resources of many institutes.

Here, we propose a mixed fast random forest (PIXANT) algorithm for multi-phenotype imputation that optimizes large data analysis in terms of runtime and memory usage by constructing fast random forest trees, making it scalable to large data of over a million individuals. PIXANT models nonlinear and linear effects across multiple phenotypes and higher-order interactions between predictive factors, allowing it to produce unbiased imputation with much higher accuracy than the aforementioned methods. We validated our method with extensive in silico simulations and compared it with the existing state-of-the-art methods: linear mixed model (LMM)^14–16^, MICE^10^, missForest^11^, and PHENIX^9^. We demonstrate that our method outperforms the competing methods in accuracy and computational efficiency. Using the UKB dataset as an example, we showed that PIXANT is orders of magnitude faster than PHENIX, with more scalable memory usage (e.g., when the sample size is 20,000 and the phenotype number is 30, PIXANT is ∼24.45 times faster and uses only about one ten thousandth of the memory compared with PHENIX). The advantages of PIXANT over PHENIX became more pronounced as the sample size increased. We then used PIXANT to impute 425 UKB phenotypes and performed GWAS. We found that more completed UKB phenotypes enabled the identification of more GWAS loci, some of which were unknown due to missing phenotypes.

## Results

### Overview of the method

PIXANT relies on the correlation between the phenotype to be imputed and other available phenotypes. PIXANT first selects reference phenotypes that are correlated with the imputed phenotype as predictors, then trains an imputation model by incorporating the linear (**b**) and nonlinear (***u***) effects among multiple phenotypes. PIXANT then imputes missing phenotypic values by projecting observable reference phenotypes into the imputation model. A flow chart is presented in **Fig. 1**. To speed up the runtime and reduce memory usage, PIXANT (1) uses a computationally efficient mix fast random forest (RF) to estimate parameters of LMM; (2) uses the Knuth^17^ (p. 137) algorithm for bootstrap resampling without replacement to construct trees for fast random forest prediction^18^; and (3) stores node information in a simple data structure, avoiding repeated accesses to the raw data and releasing memory early for efficient storage (**Online Methods**).

**Fig. 1.**
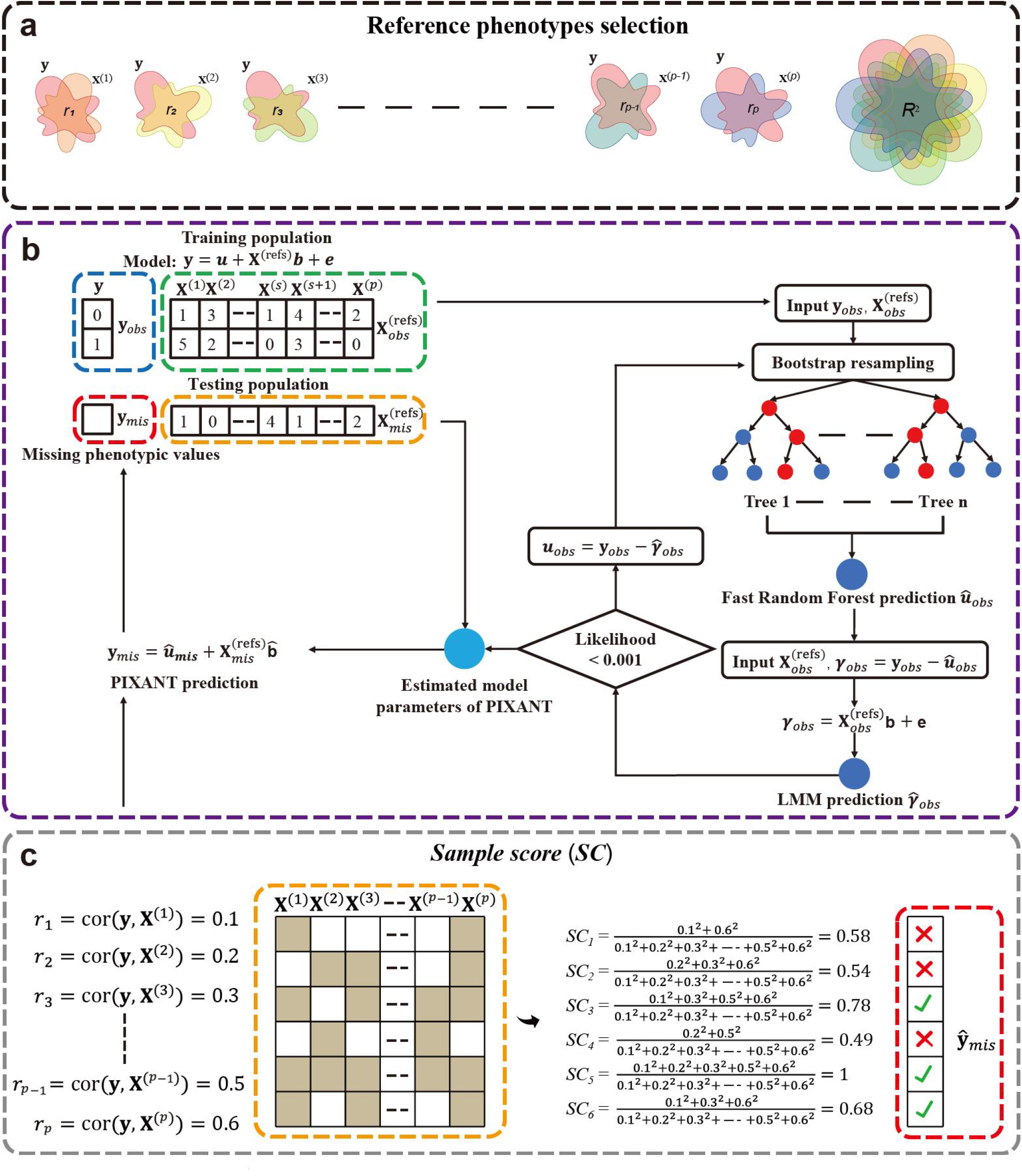
PIXANT flowchart. (**a**) PIXANT first selects *p* reference phenotypes (**X**^(refs)^) based on Pearson’s *r* at a predefined threshold using individuals overlapping with imputed phenotype **y**; then uses multiple correlation coefficient *R*^2^ of multiple regression to primarily evaluate the imputation accuracy. (**b**) After selecting the reference phenotype, PIXANT trained the model using the nonmissing imputation phenotype **y**_*obs*_ and the corresponding reference phenotypes 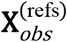. The linear effect **b** and nonlinear effect ***u*** are fitted, and the model parameters are iteratively solved by mixed fast random forest algorithm (detailed see text for details). The missing phenotypes **y**_*mis*_ is then predicted using the estimated parameters and the corresponding reference phenotype 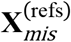. (**c**) PIXANT also performs a QC for each imputed phenotype value. In this example, assuming 6 individuals, the Pearson’s *r* between the imputed phenotype **y** and the *p* reference phenotypes are 0.1, 0.2, …, 0.6. In each individual (row), the grey grid indicates that the reference phenotype is observable and white indicates that the reference phenotype is missing, then *SC* is defined as the sum of squares of the *r* of the known reference phenotypes and the sum of squares of the *r* of all reference phenotypes. The higher the *SC* is, the more information the reference phenotype provides, and the imputed phenotype value of *SC* > 0.6 will be retained for subsequent GWAS.

For PIXANT, reference phenotypes are crucial in determining the accuracy of imputation. *R*^2^ is a metric used in training to trace the accuracy of the imputation model (i.e., how large the sample size should be to achieve an accurate imputation model). The sample score (*SC*) is defined as an individual reference phenotype missing rate that surrogates the imputation quality. Despite imputation model accuracy, the *SC* (see **Online Methods**) determines the final imputation quality of an individual (**Fig. 1**). In the following sections, we will consequently investigate how the key factors, such as reference phenotype quantity and quality, the sample size, and the *SC*, influence the performance of PIXANT in both simulations and the UKB samples.

### Computer simulations

We first investigated the performance of PIXANT in imputing missing data. A multi-trait mixed model was employed to simulate phenotypic values, which fully considered genetic and residual correlations between traits as well as relatedness between individuals to ensure phenotypic correlations (see ref.^9^ for details). We established a set of default parameters in the simulation: a sample size of 500, a heritability^9,19^ of 0.6, a number of phenotypes of 60, a coefficient of relationship between individuals^9,19^ of 0.02, and a phenotypic missing rate of 5%. We performed four simulation scenarios to investigate the performance of PIXANT: (i) sample size, 100, 300, 500, 700, 900, and 1,100, respectively; (ii) heritability, 0.1, 0.2, 0.4, 0.6, 0.8, and 1, respectively; (iii) number of reference phenotypes, 20, 40, 60, 80, 100 and 120, respectively; and (iv) relatedness between individuals, 0.02, 0.2, 0.4, 0.6, 0.8 and 0.9, respectively. We applied PIXANT and other benchmarked methods such as LMM^14–16^, MICE^10^, missForest^11^, and PHENIX^9^ in the simulations **(Supplementary Tab. 1)**. The average prediction accuracy (Pearson’s correlation) of 100 repeated experiments was used to evaluate the imputation accuracy.

As shown in **Fig. 2a**, the imputation accuracy increased with the sample size for all methods (except LMM); however, PIXANT and PHENIX outperformed all other methods over the full range of sample sizes. When the sample size was greater than 300, the imputation accuracy of PIXANT was the highest among these methods, followed by PHENIX; both had higher imputation accuracy than missForest, MICE, and LMM (**Fig. 2a** and **Supplementary Tab. 2**). For sample sizes less than 300, PHENIX achieved slightly higher imputation accuracy than PIXANT. A sample size of 1,000 appeared to deliver sufficient accuracy, where as a larger sample size contributed little to improving accuracy (**Fig. 2a**).

**Fig. 2.**
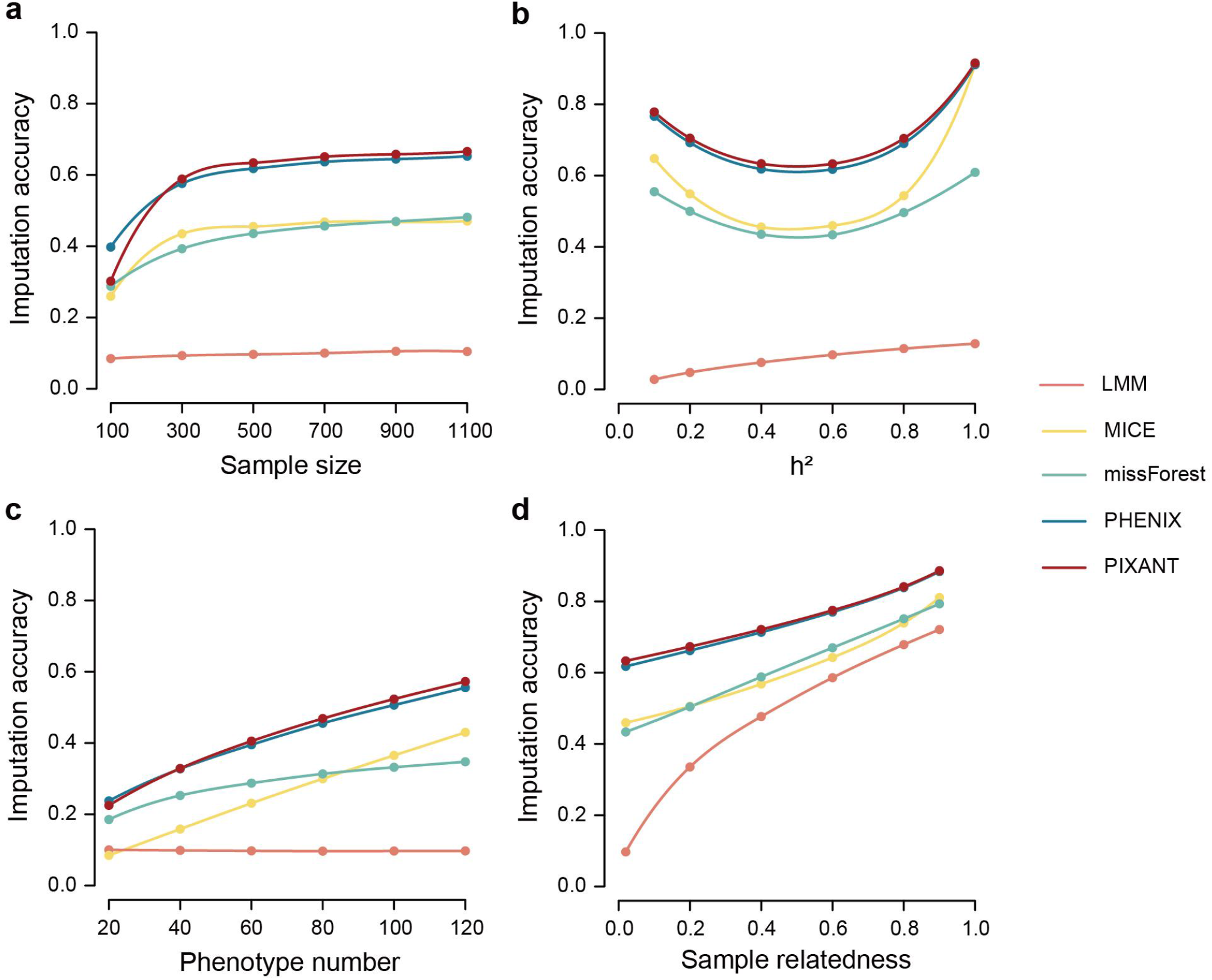
Simulation of imputation accuracy for different methods with variable sample size (a), heritability (b), number of phenotypes (c), and sample relatedness (d). Default simulation parameters were as follows: sample size of 500; heritability of 0.6; phenotype number of 60; and relationship coefficient between individuals of 0.02. Five percent of the phenotype values were set as missing before imputation. Five different methods were applied to impute the missing values. The imputation accuracy, defined as the correlation between the imputed values and the true values, is plotted on the y-axis for each method. PIXANT and PHENIX have the highest imputation accuracy in all simulations. PIXANT is slightly superior or comparable to the next most accurate method, PHENIX, except when the sample size is less than 300.

When heritability ranged from 0.1 to 1, the imputation accuracies of PIXANT were slightly higher than those of PHENIX, both of which were higher than MICE, missForest, and LMM (**Fig. 2b** and **Supplementary Tab. 3**). Increasing the reference phenotype number increased the accuracy of PIXANT, PHENIX, MICE and missForest, and the difference between PIXANT and the next best method (PHENIX) increased slightly (**Fig. 2c** and **Supplementary Tab. 4**). PIXANT and PHENIX had the highest imputation accuracy for all reference phenotype ranges and both were more accurate than other methods (**Fig. 2c** and **Supplementary Tab. 4**). The imputation accuracy of LMM was generally low regardless of the number of reference phenotypes (varying from 0.1002 to 0.0973 with the number of phenotypes *p* ranging from 20 to 120); while the imputation accuracy of PIXANT was 0.2251 for *p* = 20 and then increased to 0.5726 when *p* = 120.

LMM performance was the worst overall, except when there was a high level of relatedness between samples. When the relatedness between samples increased, the imputation accuracies of all methods improved, particularly for LMM, which explicitly relies on the relatedness between samples via the kinship matrix^20^ (**Fig. 2d**). For example, the imputation accuracy of LMM was 0.0971 when the relatedness between individuals was 0.02, but increased to 0.7211 when the relatedness between individuals was 0.9 (**Supplementary Tab. 5**).

In summary, we investigated the effects of the training model sample size and found that the genetic architecture of a typical polygenic trait of certain heritability, the number of reference traits, and levels of relatedness between samples generally influenced performance. PIXANT and PHENIX were the best methods overall in the simulated scenarios, with the former slightly better than the latter. Of note, a high kinship scenario is uncommon in population-based data sets, which often are comprised of unrelated individuals similar to the UKB dataset.

### Computational efficiency in large data

We employed UKB data^4^ comprising 277,301 unrelated White British citizens to evaluate the resource requirements of PIXANT and benchmark it against PHENIX (**Online Methods**). Note that we did not compare PIXANT with other methods that were either less accurate (LMM and MICE) or computationally prohibitive (missForest). After randomly sampling UKB subgroups of individuals (*n* ranged from 1,000 to 20,000 and *p=* 30), we evaluated the computational efficiency of PIXANT and PHENIX on a computing platform with 125 GB memory and 8 CPU cores. As shown in **Fig. 3a**, PIXANT imputation (0.06∼7.38 hours) was 0.5 to 24.45 times the efficiency of PHENIX (0.04∼0.29 hours) when performing imputation with *n* ranging from 1,000 to 20,000. PIXANT runtime was minimally affected by the sample size, whereas PHENIX runtime increased substantially with increasing sample size (**Fig. 3a** and **Supplementary Tab. 6**). The actual PIXANT memory usage was generally stable over increasing sample size (2.60 ∼ 5.12 MB for *n* ranging from 1,000 to 20,000), but PHENIX memory usage was not. For example, PHENIX required 670.49 MB memory for *n* = 1,000, but the memory usage increased to 8,052.91 MB when *n* = 20,000 (**Fig. 3b** and **Supplementary Tab. 6**). We also observed PIXANT runtime actually increased linearly with sample size and memory usage (**Fig. 3a,b** and **Extended Data Fig. 1a,b**), suggesting that PIXANT was, in principle, scalable for an even larger sample size.

**Fig. 3.**
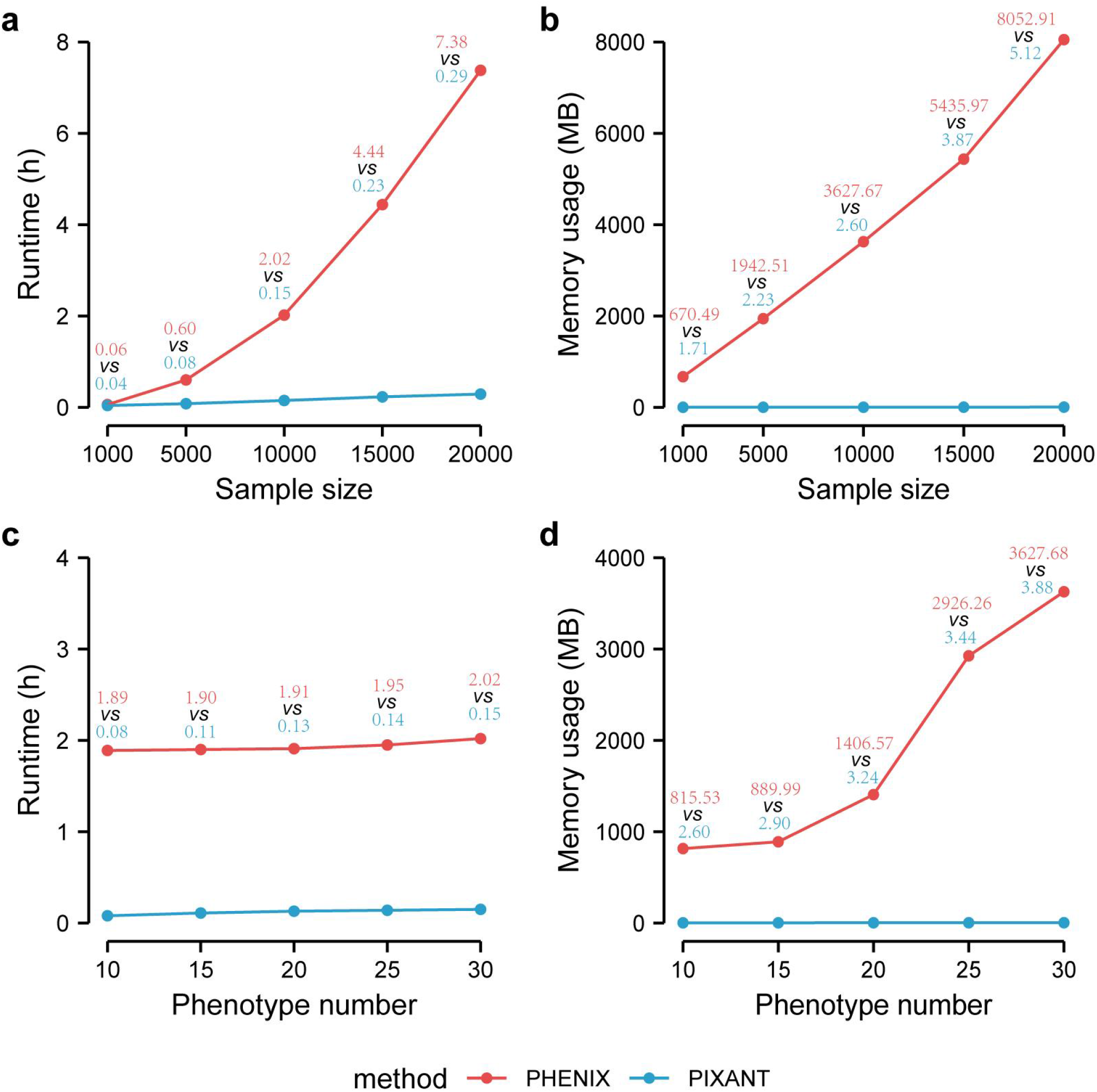
Comparison of PHENIX and PIXANT runtime and memory usage in a large sample. The effect of sample size on runtime (**a**) and memory usage (**b**), and the effect of phenotype number on runtime (**c**) and memory usage (**d**). All tests were performed in the same computing environment: 125 GB memory and 8 CPU cores. Each test was repeated an average of 5 times. The data used in the comparison of the sample size resource requirements consisted of subgroups of individuals (*n* ranged from 1,000 to 20,000 and *p* = 30) from the UKB dataset. To evaluate the resource requirements for phenotype number, we randomly sampled subgroups of individuals (*p* ranging from 10 to 30 and *n =* 10,000) from the UKB dataset. For large datasets, PIXANT is significantly less demanding on computer resources than that of PHENIX.

To evaluate the resource requirements for phenotype number, we randomly sampled subgroups of individuals (*p* ranging from 10 to 30 and *n =* 10,000) from the UKB. When the phenotype number ranged from 10 to 30, PHENIX required 1.89 to 2.02 hours, while PIXANT only required 0.08 to 0.15 hours, saving 12.47 to 22.63 times the runtime. PIXANT runtime increased slowly with variable numbers of phenotypes, as was PHENIX (**Fig. 3c** and **Supplementary Tab. 7**). Benefiting from the memory-optimized design, the actual memory usage of PIXANT was not sensitive to the number of phenotypes (∼3 MB for *p* ranging from 10 to 30), saving 312.67 to 933.97 times the computer memory compared to PHENIX (815.53∼3627.68 MB for a *p* ranging from 10 to 30) (**Fig. 3d**). We also observed that the runtime and memory usage of PIXANT increased linearly with the number of phenotypes, which allows it to process hundreds of phenotypes (**Fig. 3d, Extended Data Fig. 1c,d** and **Supplementary Tab. 7)**.

### Application in UKB data

We conducted an imputation in the 277,301 participants of the UKB dataset for 387 continuous and 38 binary phenotypes that had missing rates less than 95% and were likely to be genetically relevant (**Fig. 4a, Supplementary Data 1**). The average missing rate of these phenotypes was 21.3%. The missForest and PHENIX methods were excluded due to their prohibitive computational cost for large data, and thus only MICE was a compared. To reduce the impact of phenotypic stratification, the residual was used for imputation after correcting the phenotypes using gender, age, and the first 20 principal components that were generated from the genotypes (**Online methods**). We then selected reference phenotypes that were highly correlated (*r* > 0.3) with the imputed phenotype or the 30 most correlated phenotypes with the imputed phenotype, and obtained the number of reference phenotypes for each of the 425 imputed phenotypes ranging from 2 to 29 (median = 18). The correlation of the imputed phenotype with the reference phenotypes ranged from 0.06 to 0.99 (**Supplementary Data 1**). The correlation between the imputed and true values for 800 randomly sampled individuals was used to evaluate the imputation accuracy for each phenotype.

**Fig. 4.**
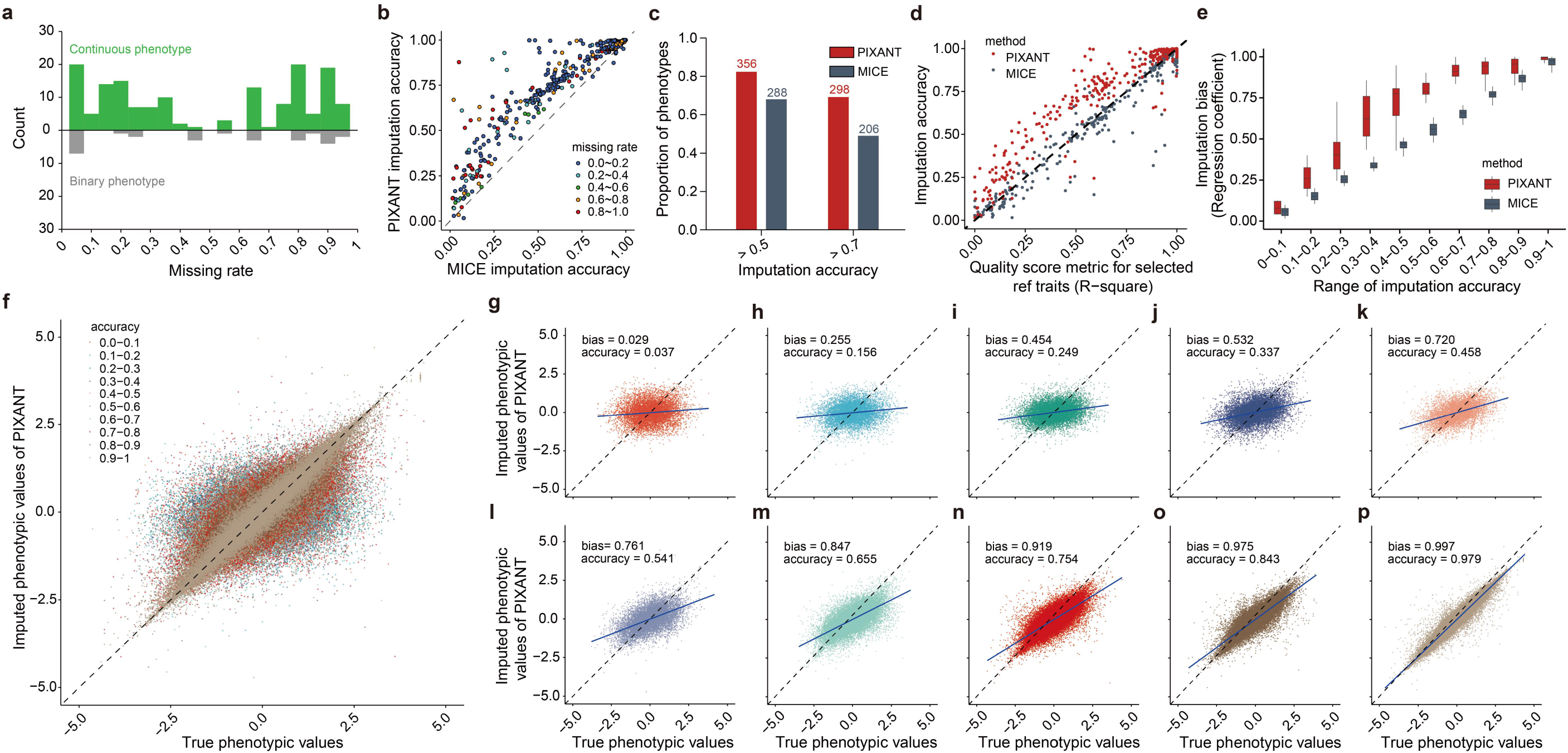
Comparison of PIXANT and MICE performance in imputation of 425 UKB phenotypes. (**a**) Distribution of phenotypic missing rates. (**b**) The imputation accuracies of PIXANT and MICE at different phenotypic missing rates, showing that the imputation accuracy of PIXANT for each UKB phenotype is generally higher than that of MICE. The phenotype missing rate is represented by different colors, with no obvious effect on imputation accuracy. (**c**) The number of accurately imputed phenotypes by PIXANT and MICE, with imputation accuracy greater than 0.5 and 0.7, respectively, showing more quality imputed phenotypes by PIXANT than MICE. (**d**) The relationship between the quality score metric *R*^2^ of the selected reference phenotypes and the imputation accuracies. The imputation accuracy of MICE is almost linear to *R*^2^, while PIXANT is a trajectory curve, implying a higher accuracy of PIXANT than MICE due to the nonlinearity property of PIXANT. (**e**),Comparison of the imputation bias of PIXANT and MICE, the imputation bias of PIXANT is better (closer to 1) than that of MICE at different ranges of imputation accuracies. (**f-p**) The imputation bias of PIXANT is shown as the observed phenotypic values (x-axis) versus imputed phenotypic values (y-axis) across all imputation accuracy ranges (**f**) and in ranges 0-0.1,0.2-0.3, etc., up to 0.9-1. (**g-p**) showing that imputation bias is improved with the increased imputation accuracy.

As shown in **Fig. 4b**, of the 425 phenotypes, 356 (83.8%) and 298 (70.1%) had PIXANT imputation accuracies greater than 0.5 and 0.7, respectively, while only 288 (67.8%) and 208 (48.9%) had in MICE (**Fig. 4c**). The relationship between the missing rate and imputation accuracy was limited (**Fig. 4b, Extended Data Fig. 2**), with a correlation of −0.176 and −0.175 for PIXANT and MICE, with *p*-values of 0.00029 and 0.00028, respectively. For example, when the missing rate of “diastolic brachial blood pressure during PWA” (Field ID: 12675) was 92.11%, the imputation accuracies of PIXANT and MICE still reached 0.947 and 0.891, respectively (**Supplementary Data 1**). The individuals with high-quality reference phenotypes (19,743 individuals in the above case) allowed for imputation despite the substantial proportion of missing phenotypes because the UKB dataset was large. Consistent with our simulation study, sample sizes greater than 1,000 were sufficient to train a good imputation model.

The imputation accuracies of the methods largely depended on the quality of the reference phenotypes. We measured the quality of the reference phenotypes with multiple correlation coefficients in multiple regression, *R*^2^, defined as the proportion of the variance explained by multiple reference phenotypes to the total variance. As shown in **Fig. 4d**, imputation accuracy increased with *R*^2^, indicating that, as expected, the quality of the reference phenotypes played an important role in determining accurate imputation. Additionally, the imputation accuracy of MICE was nearly linear to*R*^2^, while the accuracy of PIXANT was a trajectory curve, implying its advantage in drawing nonlinear information from the reference phenotypes (**Fig. 4d)**.

We found that the PIXANT imputation bias (**Online Methods**) was better (closer to 1) than that of MICE for different ranges of imputation accuracies (**Fig. 4e**), suggesting that imputation with PIXANT led to more reliable GWAS signals than that with MICE. Furthermore, the imputation bias was correlated with the imputation accuracy; as the imputation accuracy increased, the imputation bias improved for both methods (**Fig. 4f-p**). For example, when the imputation accuracy was between 0 and 0.1, the imputation bias of PIXANT was 0.029 (**Fig. 4g**), but when the imputation accuracy was between 0.9 and 1, the imputation bias of PIXANT increased to 0.997 (**Fig. 4p**), suggesting that a higher accuracy could produce a more unbiased imputation.

We next ask if the imputation is statistically significant. To this aim, we randomly permuted part of the reference phenotypes in a range of proportion, respectively, we observed the imputation accuracy declined as more proportion of reference phenotypes disturbed, demonstrating the validity of the imputation. We further determined the empirical *p*-value by permuting the whole reference phenotype randomly. We selected 12 phenotypes randomly with imputation accuracy between 0.025 and 0.333, and we expected high accuracy would results in significant imputation. Indeed, when the imputation accuracy was greater than 0.1, the imputation in UKB dataset will be significant (**Extended Data Fig. 3**).

### Individual-level quality control

Even when with a phenotype imputation accuracy greater than 0.5 or 0.7, some individuals of the phenotype may not be well imputed if an individual’s correlated reference phenotype records are also missing. For example, GWAS of the QT interval phenotype (the time between the start of the Q wave and the end of the T wave in the heart’s electrical cycle, Field ID: 22331, imputation accuracy = 0.671) identified far fewer loci after imputation with PIXANT than without imputation (**Fig. 5a**). Because most individual imputed phenotypes had few or no reference phenotype records, we developed a *SC* for quality control of an individual imputed phenotype that quantifies how one’s nonmissing reference phenotypes contribute to an imputation task (**Online Methods**). The *SC* range is 0∼1, and a higher *SC* can lead to more precise phenotype imputation. For the QT interval phenotype, we filtered the imputed individual phenotypic values with different *SC* thresholds ranging from 0 (no reference phenotypic values) to 1 (complete reference phenotypic values) and evaluated the imputation accuracy at different *SC* levels using 800 phenotypic values (as previously described). We found that 78.2% of the imputed phenotypes had *SC* less than 0.2 (**Fig. 5b**), which explained the low power of GWAS after imputation of the QT interval phenotype. We further examined if the developed *SC* was able to well approximate the real imputation accuracy at the individual level. We determined the imputation accuracy and *SC* for these 356 phenotypes (with an imputation accuracy greater than 0.5) and observed that the imputation accuracy increased proportionally to the *SC*, well approximating the imputation accuracy for each individual (**Fig. 5c**). We found that the imputation accuracy increased sharply between 0 and 0.2 but gradually from 0.2 to 1 (**Fig. 5c**), suggesting that *SC* greater than 0.2 might be a practical threshold that could lead to an accuracy greater than 0.7.

**Fig. 5.**
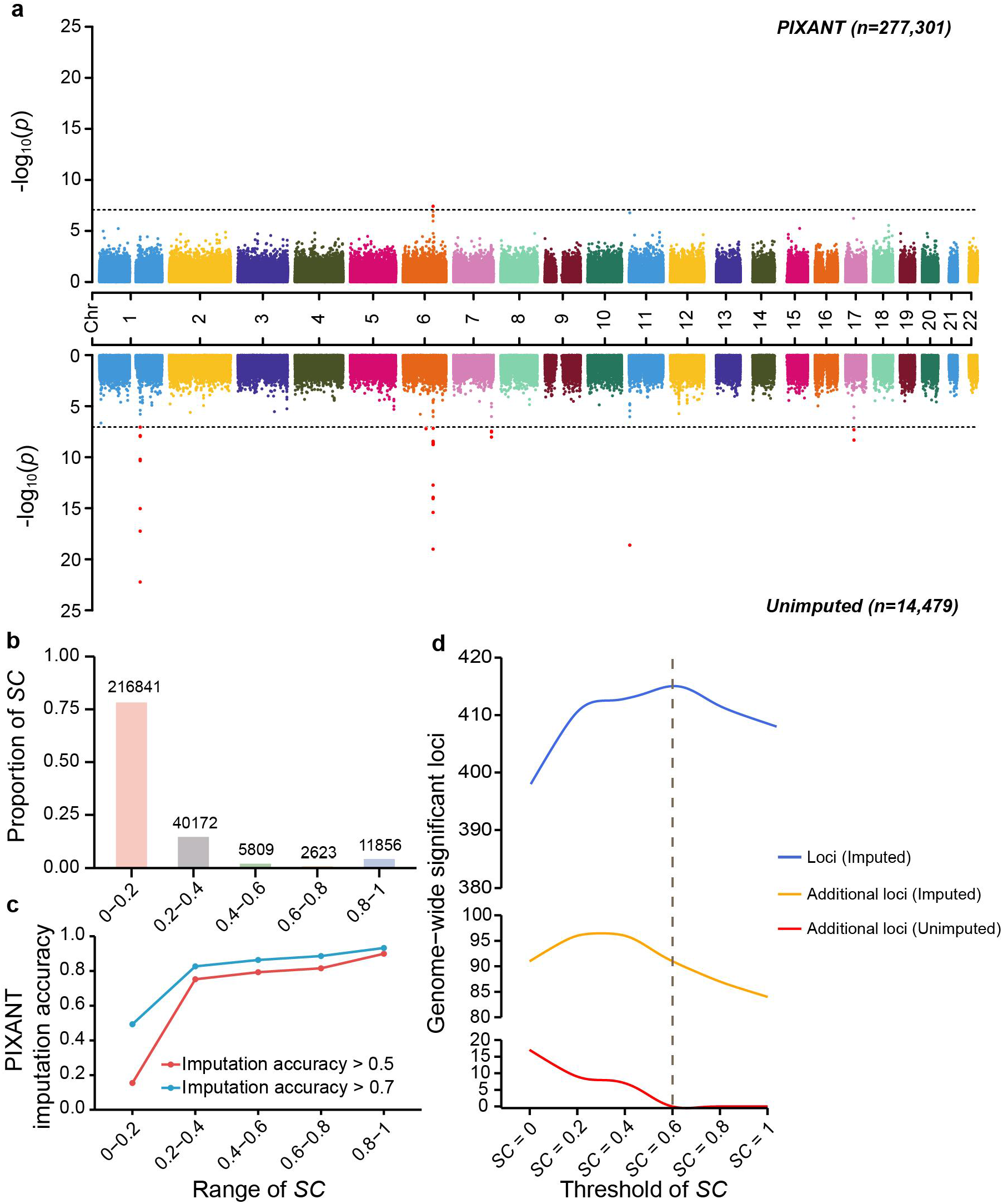
The relationship between *SC*, power, and PIXANT imputation accuracy in the UKB dataset. (**a**) GWAS of the UKB QT interval phenotype with imputed (upper panel) and unimputed (lower panel) phenotypes, showing that the GWAS signals are attenuated due to low imputation quality of some individuals, which prompted us to develop *SC* to implement a QC for each individual. (**b**) The frequency of imputed QT interval phenotypes under different *SC* thresholds, more than 75% of individual phenotypes are imputed with an *SC* of less than 0.2, which explained the attenuated GWAS signals after imputation. (**c**) The imputation accuracy across 42 randomly selected UKB phenotypes under different *SC* thresholds. *SC* is in good agreement with the true imputation accuracy, implying *SC* is a reasonable QC at the individual level. (**d**) GWAS power for 42 randomly selected phenotypes before and after phenotype imputation under different *SC* thresholds. The total number of GWAS loci after imputation (upper panel), the additional GWAS loci after imputation (middle panel), and the additional GWAS loci before imputation (bottom panel). The latter reflects the GWAS loci overlap before and after imputation, fewer numbers imply a higher overlap. *SC* of 0.6 is the optimal threshold, leading to the highest power and the maximum overlap, while *SC* between 0.2-0.6 also seems to offer sufficient power and overlap.

To refine the *SC* threshold, we randomly selected 42 imputed phenotypes, filtered individual phenotypic values with thresholds of 0, 0.2, 0.4, 0.6, 0.8, and 1, and performed the GWAS with the filtered phenotypes. An *SC* of 0.6 (i.e., an imputation accuracy of approximately 0.8) optimized the GWAS power (upper panel in **Fig. 5d**) and resulted in the largest GWAS loci overlap before and after imputation (bottom panel in **Fig. 5d**). The GWAS power and GWAS loci overlap was stable when *SC* was between 0.2 and 0.6 (upper panel of **Fig. 5d**), which is thus recommended for individual-level imputation filtering.

### UKB GWAS after PIXANT imputation

We performed GWAS of the 425 imputed UKB phenotypes to evaluate the usefulness of our method, positing that a more effective imputation would identify more GWAS associations. We retained only phenotypes with imputation accuracy greater than 0.5 or 0.7 (356 and 298 phenotypes, respectively) to ensure the GWAS reliability (**Supplementary Data 2**). After removing individual phenotypes with *SC <* 0.6 for each UKB phenotype, we performed GWAS with PLINK^21^ and then applied a genomic control factor for potential population stratification^22^. A conservative *p*-value threshold of 8.87× 10^-8^ after Bonferroni correction was employed to identify significant GWAS loci (**Online Methods**).

The imputation accuracy of PIXANT was 0.772 for heart rate (Field ID: 5983), which had a missing rate of 84.92%. We compared the GWAS heart rate results using imputed (*n* = 271,860) and unimputed (*n* = 40,990) phenotypes (**Fig. 6a**). Six independent GWAS loci (>1 Mb from each other) were identified before imputation. The GWAS signal was increased after imputation, and another 31 independent loci were identified (**Fig. 6a** and **Supplementary Tab. 8**). The plot highlights several peaks containing genes related to dilated cardiomyopathy, cardiac rhythm regulation, the cardiac parasympathetic nervous system, heart development, and heart disease (*DSP* (ref.^23,24,25^), *CDH11* (ref.^26,27^), *RNF220* (ref.^28,29^), *SCN10A* (ref.^30,31^), *MYH11* (ref.^32^) and *RGS6* (ref.^33,34^), etc.) (For details, see **Supplementary Tab. 8**).

**Fig. 6.**
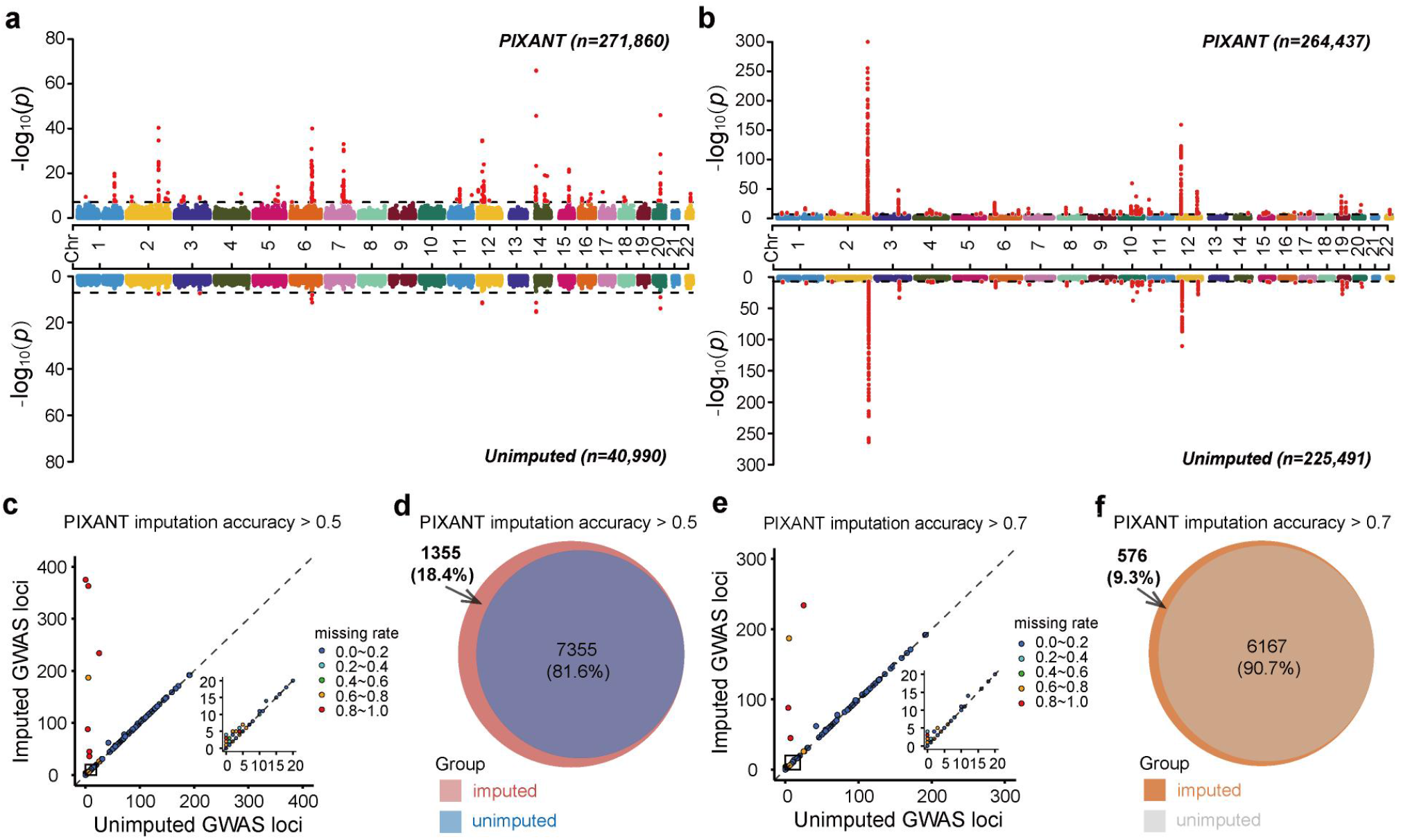
GWAS of 425 UKB phenotypes before and after imputation. (**a**) Heart rate example for imputed (upper panel) and unimputed (lower panel) phenotypes. The dashed horizontal lines denote a significance threshold of 6.74 (-log10(0.05/total SNPs)). After imputation, the GWAS signals are significantly increased and more GWAS loci are identified. (**b**) Direct bilirubin example. (**c**) Comparison of GWAS loci before and after imputation for 356 UKB phenotypes with imputation accuracy greater than 0.5. Each point represents an imputed phenotype, and different colors represent different phenotypic missing rates. The difference in the number of GWAS loci before and after imputation is more pronounced as the phenotypic missing rate increases. (**d**) Venn diagrams showing GWAS loci overlap before and after imputation of 356 phenotypes; 18.4% more GWAS loci were identified after imputation than before (8,710 *vs* 7,355). (**e**) Comparison of GWAS loci number before and after imputation for 298 UKB phenotypes with an imputation accuracy greater than 0.7. (**f**), Venn diagrams showing GWAS loci overlap before and after imputation of 298 phenotypes; 9.3% more GWAS loci were identified after imputation than before (6,743 *vs* 6,167).

The direct bilirubin phenotype (Field ID: 30660) had a missing rate of 14.73% and an imputation accuracy of 0.910. The GWAS signals were significantly enhanced after imputation, and additional GWAS loci were found on chromosomes 1, 2, 3, 4, 6, 10, 15, and 22 (**Fig. 6b** and **Supplementary Tab. 8**). Interestingly, a lead SNP within *CSP1* of chromosome 2 was associated with gallbladder disease: clinicopathological characterisation showed that increased expression of *CPS1* is associated with reduced levels of international normalized ratio (INR), total protein and alkaline phosphatase (ALP), and indirect bilirubin^35^. A lead SNP within *GSTM1* on chromosome 1, whose null genotype has a high risk of pathologic hyperbilirubinemia, may cause higher bilirubin levels^36^. In addition, *GSTM1* was also associated with total bilirubin levels in neonatal jaundice^37^ (**Supplementary Tab. 8**).

We then summarized the GWAS results for all 356 phenotypes with an imputation accuracy greater than 0.5. After imputation, the number of GWAS loci was significantly higher than before imputation (**Fig. 6c)**. In total 7,355 independent GWAS loci were identified before imputation, while 8,710 were identified after imputation, an increase of 18.4% (**Fig. 6d**). As expected, the difference in the number of GWAS loci before and after imputation became more pronounced as the phenotypic missing rate increased (**Fig. 6c)**. The results of 298 phenotypes with an imputation accuracy greater than 0.7 were similar to the above results (**Fig. 6e**): a total of 6,167 GWAS loci were identified before imputation, and 576 (9.3%) additional loci were identified after imputation (**Fig. 6f**). These findings suggest that with reasonable quality control at both the phenotype and individual levels, imputation may significantly improve GWAS power. The GWAS results are detailed in **Supplementary Data 3**.

To further confirm our findings, we tried to replicated some of the GWAS results. We first masked half of the direct bilirubin phenotype (Field ID: 30660), and imputed them with PIXANT, followed by GWAS, the GWAS results agreed well with using complete datasets (**Extended Data Fig. 4**). In addition, we compared the additional human height (Field ID: 12144) GWAS loci identified by PIXANT with GIANT summary statistics based on meta-analysis of multiple populations independent of UKB^38^, as expected, 99.4% of UKB GWAS loci were significant in GIANT despite the weaker signals in UKB, and the GWAS signals of both are well correlated (Person’s *r* of -log10(*p*) is 0.949) (**Extended Data Fig. 4**), demonstrating the validity of the imputation with PIXANT.

## Discussion

Missing genotype and phenotype data are inevitable in large-scale genetic datasets. Here, we developed a new multi-phenotype imputation method, PIXANT. The major advantage of PIXANT is its computational speed and stable memory usage during analysis of big data. When the method is applied to data from millions of individuals, PIXANT is several orders of magnitude better in runtime and computer memory usage than the state-of-the-art method PHENIX. Our simulation and real data analyses demonstrated that our method was comparable to or outperformed state-of-the-art methods for accuracy. In addition, the analysis of real data and the biologically plausible results suggest that imputation can uncover new true positive associations and plausible candidate genes (**Fig. 6a** and **Supplementary Tab. 8**). For example, we identified an additional locus with the lead variant rs272570 within *RNF220*, a gene that is strongly associated with heart rate. This association has also been reported in previous studies^28,29^.

Like many multi-phenotype imputation methods^10,39^, combining high-quality reference phenotypes can improve the imputation accuracy of PIXANT. While PIXANT successfully imputed most of the phenotypes in the UKB dataset, it was unable to impute missing phenotypes when few or no highly correlated phenotypes were recorded. This suggests that the reference phenotypes selected for imputing must be highly relevant. Similar to existing methods, the imputation accuracy of PIXANT is influenced by the quality of reference phenotypes involved, meaning that a comprehensive phenotype dataset is needed to provide highly correlated reference phenotypes for PIXANT. In this study, we defined the quality score of the selected reference phenotype as the multiple regression *R*^2^ between the imputed phenotype and its reference phenotypes, which is primarily useful in evaluating the imputation accuracy of the selected reference phenotypes. Our results also suggest that only high-quality imputation will enhance signals; poor imputation for some individuals may attenuate the association signals. We consequently propose an individual-level QC metric *SC* for PIXANT. We found that applying *SC* was essential for improving power and controlling false-positive signals in GWAS. Our real data analysis demonstrated that the *SC* threshold was 0.2-0.6 (**Fig. 5**). We recommend an *SC* threshold of 0.6 to output high-quality imputed phenotypes.

PIXANT allowed a high missing rate in the UKB data imputation. A substantial portion of phenotypes were missing in the UKB dataset, but because the dataset was large, the existing high-quality observable phenotypes allowed for reliable imputation. Consistent with our simulation study, a sample size greater than a certain number (typically tens of thousands in the case of UKB data) is sufficient to train a good imputation model (**Fig. 2a** and **Supplementary Tab. 1**). A large number of binary phenotypes are recorded in the UKB data in addition to continuous phenotypes. In this study, we directly treat a binary trait (0/1) as a continuous for imputation, and it usually performs well in practice. The rationale behind this method is that it can leverage the full range of statistical techniques available for continuous data, which may not be fully utilized when the data are strictly dichotomized. The current version of PIXANT is not applicable for imputation of categorical traits, but it can be adapted to accommodate such scenarios. The core of PIXANT is the random forest, which can solve binary and categorical traits using classification trees rather than regression trees for continuous phenotypes, thus warranting further study.

PIXANT can be applied in the imputation of multi-omics data, such as those in the GTEx project datasets. The GTEx project contains nearly ∼10 thousand RNA-seq records from 44 tissues^40–42^ and plays an important role in the identification of functional genes that underlie human diseases^43,44^. For some tissues, only a subset of individuals is sequenced, resulting in missing data in the gene expression matrix. PIXANT is able to impute the “missing” gene expression data by leveraging gene expression data from other tissues, using the overlapping sequenced individuals as a reference panel. When the number of tissues is large, gene expression correlations between tissues are widespread, providing more information for imputation. As high-dimensional imputation will be computationally intensive, current imputation algorithms are extremely challenging to employ at such a scale. PIXANT may offer an alternative for multi-omics imputation. In the future, PIXANT may be applied to omics data, such as chromatin immunoprecipitation following by sequencing (ChIP-seq)^45^, assay for transposase-accessible chromatin with sequencing (ATAC-seq)^46^, and whole genome bisulfite sequencing (WGBS)^47,48^. Imputation for high-dimensional multi-omics data will be favorable for discovering genomic regulatory elements and will become more common for further exploring causal genes and pathways underlying human traits and diseases. However, it is not trivial to achieve high computational efficiency for imputation, which will be central to further development of the present work.

## URLs

UK Biobank, http://www.ukbiobank.ac.uk;

PLINK (v1.90 beta), https://www.cog-genomics.org/plink2;

MissForest (v1.5.0), https://cran.r-project.org/web/packages/missForest/index.html;

MICE (v3.16.0), https://cran.r-project.org/web/packages/mice/index.html;

PHENIX (v1.0), https://mathgen.stats.ox.ac.uk/genetics_software/phenix/phenix.html;

PIXANT (v0.1.0), https://github.com/GuLinLin/PIXANT;

GIANT summary statistics: https://portals.broadinstitute.org/collaboration/giant/images/4/40/GIANT_HEIGHT_YENGO_2022_GWAS_SUMMARY_STATS_ALL_excluding_UKB.gz.

## Acknowledgements

We thank the participants of UK Biobank for making this work possible (under application 41376). The research was supported by the National Key Research & Development Program of China (2022YFD2401001); National Natural Science Foundation of China (31771392), the Natural Science Foundation of Fujian Province of China (2021J02045); the Seed Industry Innovation and Industrialization Project of Fujian Province (2021FJSCZY01); the National Marine Fisheries Industrial Technology System Post Scientist Project ([CARS-47-G04](https://www.sciencedirect.com/science/article/pii/S0044848623004568)).

## Author contributions

FM, JD, and CGB conceived and supervised the study. GLL conceived and developed the method, wrote the program and conducted the whole analysis. WHS, LTY and ZYJ assisted in the data collection and contributed to the demonstration of the results. CGB and JCH contributed to real data analysis and conceived the imputation quality control method. FM, CGB, LXL and WZY discussed results and methods, and provided comments that improved earlier versions of the manuscript. FM, JD, and CGB contributed resources and funding. GLL, FM and JD drafted the manuscript, and CGB revised the first draft of the manuscript. All authors reviewed and approved the final manuscript.

## Data availability

The data in this manuscript from the UK Biobank (http://www.ukbiobank.ac.uk) was available dependent on successful application the data access committees. This research has been conducted using the UK Biobank Resource under Application Number 41376.

## Online methods

### Multi-<u>p</u>henotype <u>i</u>mputation with mi<u>x</u>ed f<u>a</u>st ra<u>n</u>dom fores<u>t</u> (PIXANT)

PIXANT fits linear effects and nonlinear (high-order interactions) of *p* reference phenotypes. Assuming that *n* individuals are considered, the model can be written as,

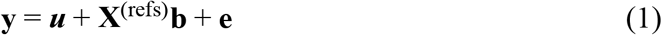

where **y** is a *n* × 1 vector of imputed phenotypic values, **X**^(refs)^ is a *n* × *p* matrix, denotes *p* reference phenotypic values; ***u*** is a *n* × 1 vector, denoting the nonlinear effect vector for *p* the reference phenotypes selected for imputation, **b** is a *p* × 1 vector, denoting the linear effect vector for the *p* reference phenotypes; **e** is *n* × 1 vector, denoting the random error, which follows a normal distribution.

Before imputation, PIXANT first trains the predictive model using observable phenotypes. We denote the observed phenotypic values by **y**_*obs*_ and the missing phenotypic values by **y**_*mis*_ ; and similarly, we use 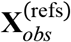 denote *p* reference phenotypic values correspongding to **y**_*obs*_ individuals, and 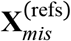 to denote reference phenotypic values corresponding to **y**_*mis*_ individuals. Then, the training model can be expressed as

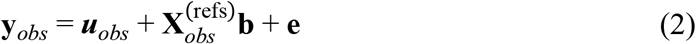

We solve the model parameters of equation (2) using an iterative algorithm. Before updating, we first sorted reference phenotypes according to the missing rate from low to high.

1. *Initialization*. First, we initialize the model parameters with **b =** 0 and ***u* = y**. **X** ^refs^ is initialized with average observed phenotypic values or randomly sampled from the nonmissing values ^11^.
2. *Update of linear-effect parameters*. Let

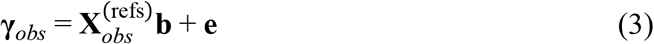

representing the phenotypic values attributed to the linear effect and random error, where **γ**_*obs*_ at *t*^th^ iteration equals to **γ**_*obs t*_ = **y**_*obs*_ - ***u***_*obs t*-1_ by substituting equation (3) to equation (2); then **b** _*t*_ is updated by solving linear regression of equation (3).
3. *Update of nonlinear effect parameters*. Let

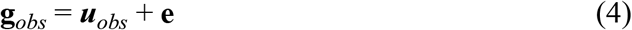

representing the phenotypic values attributed to nonlinear effect and random error. We fit a nonlinear fast random forest^49^ function 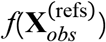 for ***u***_*obs*_, i.e.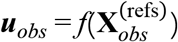, reflecting the complex nonlinear effect of p reference phenotypic values on the imputed phenotype. Then equation (4) can be written as

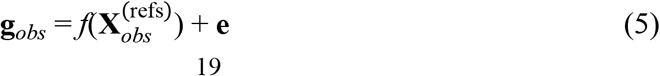

where **g**_*obs*_ at *t*^th^ iteration can be obtained by

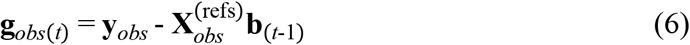

after substituting equation (4) to (2). Then the parameters involved in the fast random forest function 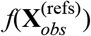at iteration *t*^th^ can be estimated by solving the random forest nonlinear regression of equation (5). To speed up the runtime and reduce the memory usage, we use a computationally efficient mix fast random forest (RF) to estimate model parameters implemented via efficient sampling and data storage techniques (see the main text for more details). We therefore have the update of the nonlinear effect ***u***_*obs t*_ with estimated parameters in 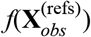.

We update steps (2)-(3) repeatedly, until the change in likelihood reaches the given criteria (< 0.001). In practical applications, the model usually converges in a few iterations and is not sensitive to the initial values.

In the same way, we also train the model for *p* reference phenotype, sequentially. The model can be written as

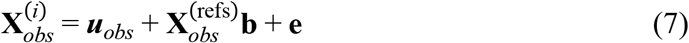

for *i* = 1, 2, …, *p*, which is similar to equation (2), except that **y**_*obs*_ is substituted by 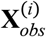, thus 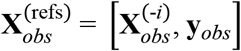. The model parameters for each reference phenotype in equation (7) are also solved using an iterative algorithm as described above.

We have completed the model training, and next we are going to impute missing phenotypic values using trained model parameters. We use the prediction model.

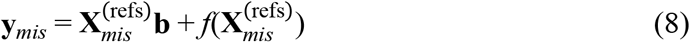

to update **y**_*mis*_, and use the predict model

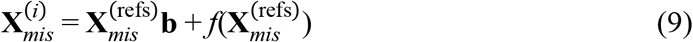

to update 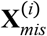, for *i* = 1, 2, …, *p*, where 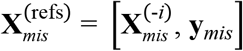. PIXANT updates **y**_*mis*_ and 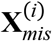 iteratively, using the most recently updates whenever possible. The update is repeated for several times until the difference of missing phenotypes 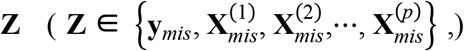 in two consecutive iterations meets a certain criterion (θ < 0.001), where θ is defined as

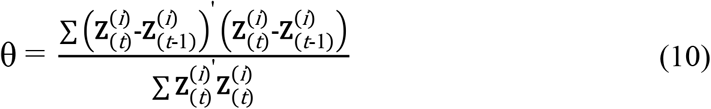

### Imputation accuracy

We assessed the imputation accuracy with the correlation coefficient between the imputed and true phenotypic values,

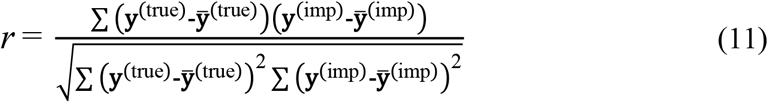

where **y** ^true^ is the true phenotypic values and **y** ^imp^ is the imputed phenotypic values. The correlation coefficient was calculated with the function “cor.test” in R package “psych” (v2.4.3).

### Sample score (*SC*)

Although we have filtered out poor imputed phenotypes, in every phenotype with good imputation, the poorly imputed individual phenotype will have a non-negligible effect on GWAS, which prompted us to develop a sample score for QC for an individual phenotype. When imputing for individual *i*, assuming it has *p* reference phenotypes, of which *p*^*’*^ phenotypic values are complete while others are missing, then *SC* is defined as

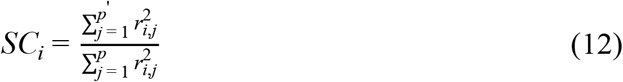

where, *r*_*i*,*j*_ is the correlation coefficient between *j*^th^ (for *j* = 1, 2,…, *p*) reference phenotype and the imputed phenotype of individual *i*. We empirically determined the optimal QC threshold of 0.2-0.6 by the number of (additionally) identified GWAS loci before and after imputation.

### Imputation bias

Although we have used imputation accuracy to assess the effect of the imputation, it reflects the correlation between imputed and real values and cannot evaluate the degree of systematic deviation between the imputed values and real values, while biased imputations may mislead the GWAS results. On the basis of this consideration, we further assessed the bias in the imputation. To do this, we randomly selected an additional 0.29% of the complete phenotypic values missing (800 phenotypic values) from the imputed phenotype, and then evaluated the imputation bias using the regression coefficient between the imputed and the true phenotypic values, where the simple regression analysis was implemented via the function “lm” in R (v4.2.0).

### PHENIX, missForest, MICE, and LMM

We applied several other popular methods for imputing missing phenotypic values (**Supplementary Tab. 1**). We herein provide a brief introduction of each method.

*PHENIX*. The PHENIX method is expressed as

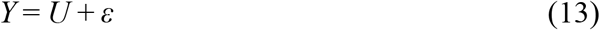

where, *Y* is a *n × p* matrix of *p* phenotypes measured on *n* individuals, which is rescaled across each phenotype, so that the mean and variance of each trait is 0 and 1, respectively; *U* and *ε* is defined as a matrix normal distribution,

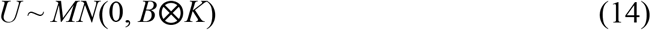

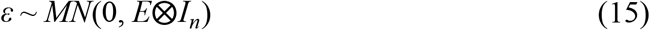

where *U* and *ε* represent *N* × *P* matrix that column-wise restores the *NP* × 1 matrix, respectively, *K* is the *n* × *n* kinship matrix between individuals, *I*_*n*_ is the *n* × *n* identity matrix; *B* and *E* is the *p* × *p* matrix, representing genetic and residual covariances between phenotypes, respectively. PHENIX uses the variational Bayes method to iteratively update missing phenotypic values *Y* ^*miss*^ by approximating joint posterior distribution of model parameters (For details, see refs^9^).

*missForest*. MissForest is a kind of multi-phenotype imputation method based on the random forest method. It first makes an initial guess of the missing phenotypic values with the mean/median of the corresponding phenotypes; then sorts the phenotypes according to the missing rate and imputes the missing values of each phenotype in order with a random forest method^11^.

*MICE*. MICE was developed by Buuren and Oudshoorn^10^, which is based on the multivariate multiple imputation scheme of Schafer (2002)^50^. Let the hypothetically complete data *Y* be a partially observed random sample from the *p* -variate multivariate distribution *Y*|*θ* with parameter *θ*, the MICE algorithm obtains the posterior distribution of *θ* by sampling iteratively from conditional distributions of

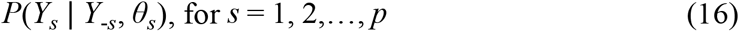

The model parameters are successively drawn with Gibbs sampling. Specifically, for *t*^th^ iteration, they are sampled from the conditional posterior probability distribution,

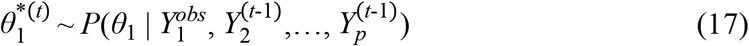

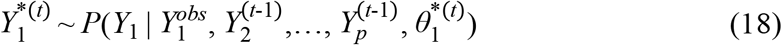

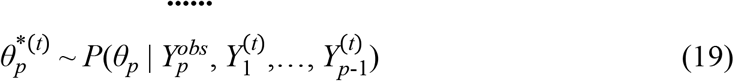

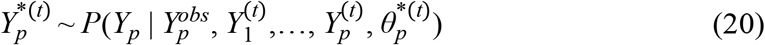

where 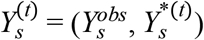 is the *s*^th^ imputed variable at iteration *t*.

*LMM*. The linear mixed model (LMM) method is a kind of single-trait method, which imputes missing phenotypic values with the BLUP technique. For each phenotype, LMM is defined as

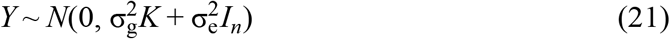

where *Y* denotes the phenotype, 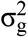 and 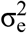 are genetic and residual variances, respectively; *K* is the kinship between individuals, which is derived from whole genome SNPs or pedigree^51^.

### Simulations

We simulated genetically and environmentally related phenotypic data with multiple phenotypes according to the method of Dahl and Baud^9^. The method dissects the phenotypic values into genetic and environmental effects. The phenotype was simulated with a model

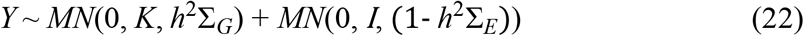

where *Y* is *n × p* the multi-trait phenotypic values that follows a multivariate normal distribution; *K* is the relationship matrix between individuals; *h*^2^ is the heritability and Σ_*G*_ is the *p × p* the genetic correlation matrix between traits, which is sampled from Wishart distribution, Wi(I_*p*_, *p*, df), where df = *p*. The small degree of freedom *p* will produce a covariance matrix with a large variation; Σ_*E*_ is the *p × p* residual covariance matrix, which is also sampled from Wishart distribution, Wi(*I*_*p*_, *p*, df) with df of *p*. The genetic and environmental effects are randomly sampled from the multivariate normal distribution *MN*(0, *K, h*^2^Σ_*G*_) and *MN*(0, *I*, (1-*h*^2^Σ_*E*_)), respectively, and the phenotypic values are simulated with equation (22). The simulation is based on Dahl and Baud’s script^9^. To make it suitable for simulations with a large unrelated population we modified the parameter “fam_size” to “sample size”. We then randomly selected a certain percentage of phenotypes as missing.

### UK Biobank data

The UK Biobank Project contains half a million UK participants, aged between 40 and 69 years at the time of recruitment. All individuals in the project were genotyped for approximately 800,000 SNPs using the Applied Biosystems UK BiLEVE Axiom Array by Affymetrix^4^. A wide variety of phenotypic and health-related information is available for each participant, including biological measurements, lifestyle indicators, biomarkers in blood and urine, and imaging of the body and brain.

### Quality control

We extracted the unrelatedness individuals from British citizens for study. After removing individuals with certain kinship (Field ID: 22006, Genetic ethnic grouping; Field ID: 22021, Genetic kinship to other participants), 278,788 unrelated individuals were retained. We then used PLINK^21^ (v1.90 beta) software for quality control (QC) to remove individuals with missing genotypes > 5% and SNPs with missing rates > 5%, Hardy-Weinberg equilibrium *p*< 1×10^-6^ and MAF < 0.01. We also checked the samples for sex discrepancies with parameter “--check-sex” and removed the samples marked as “PROBLEM”; we then removed the individuals with excess heterozygosity with parameter “--het”, and excluded the samples with the heterozygosity exceeding three standard deviations. Finally, the data included 277,301 individuals of British ancestry with 563,675 common SNPs (MAF>1%).

### Imputation of UK Biobank data

For each phenotype to be analyzed, we selected a number of reference phenotypes that are correlated to it (*r* > 0.3 or top 30 traits) for imputation. We used the function “corr.test” of the R package “psych” (v2.4.3) with the parameters use=“pairwise”, method=“spearman”, adjust=“fdr”, alpha=0.05, to calculate the correlation coefficient. We also performed principal component analysis with the genotype data, which was implemented with the function “bed_randomSVD” of the R package “bigsnpr”^52,53^. The top 20 principal components were used to construct the covariance matrix to represent the population structure. For each phenotype, we used the linear model to correct for the principal components and appropriate covariates and used residual **y’** for imputation.

### GWAS and functional annotations

The phenotypes after imputations were used for genome-wide association analysis. We used a linear model **y’** = ***μ*** + SNP + **e** for GWAS, which was implemented via PLINK^21^ (v1.90 beta). We defined the whole-genome significance cutoff as the Bonferroni test threshold, which was set as 0.05/total SNPs^54^ (α = 0.05/563,675, *p*-value < 8.87×10^-8^). We defined GWAS signals with a distance greater than 1Mb as mutually independent loci, we then summarized the number of independent loci identified in GWAS studies. Within the GWAS loci, we used the UCSC browser to annotate the functional genes in the GWAS region according to human genome assembly GRCh37, and screened the functional candidate genes in the 500kb region around the lead SNP.

**Extended Data Fig. 1.**
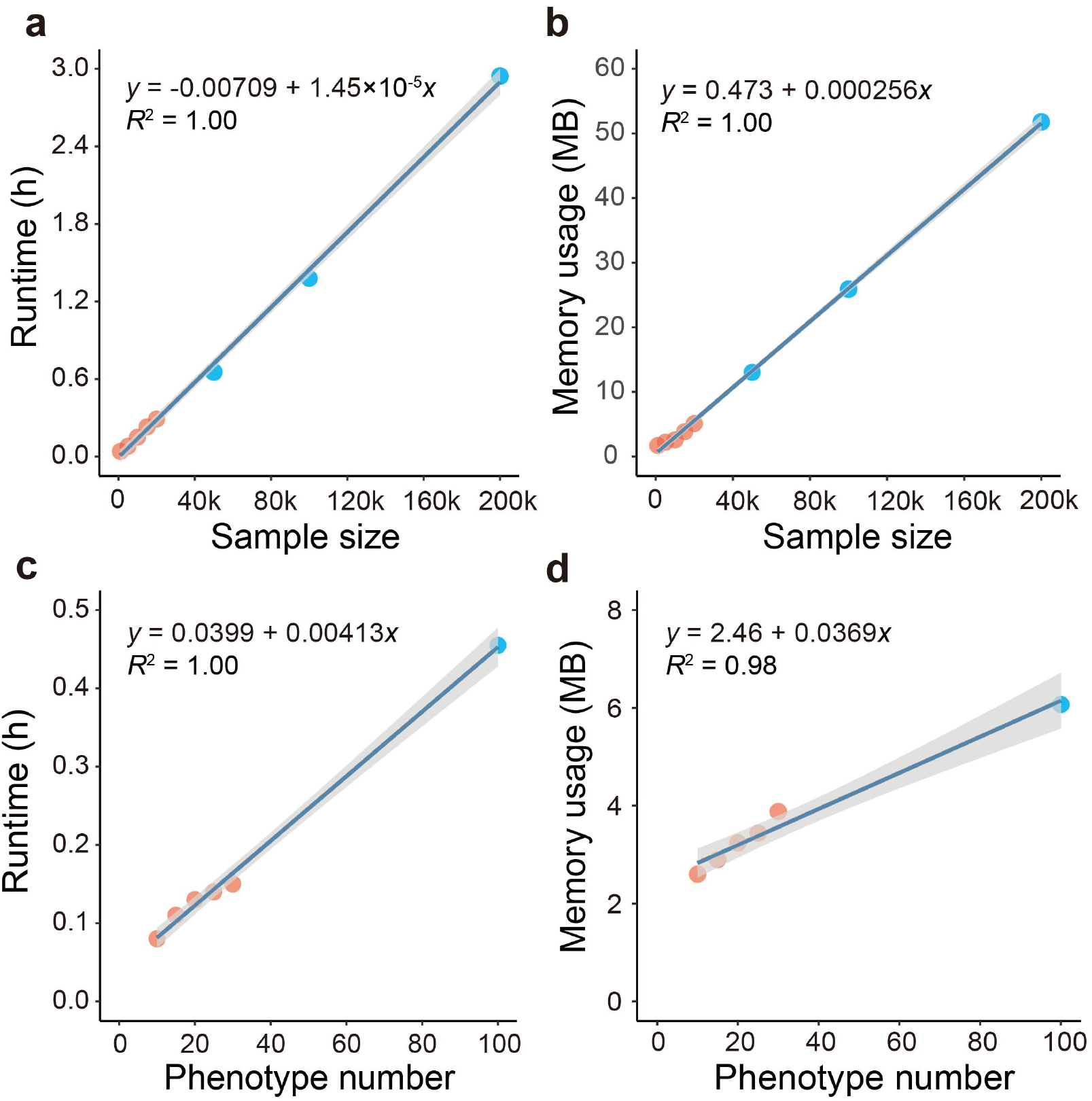
The computational resources required by PIXANT to increase sample size and number of phenotypes. (**a**) and (**b**), further increasing the sample size from a few hundred to 50k, 100k, and 200k, the runtime and memory usage increase approximately linearly. (**c**) and (**d**), As the number of phenotypes further increases from 5, 10, 15, 20, 25, and 30 to 100, the runtime and memory usage also increase approximately linearly. The results demonstrate that PIXANT scales linearly with both sample size and phenotype number, making it suitable for handling large datasets with multiple traits.

**Extended Data Fig. 2.**
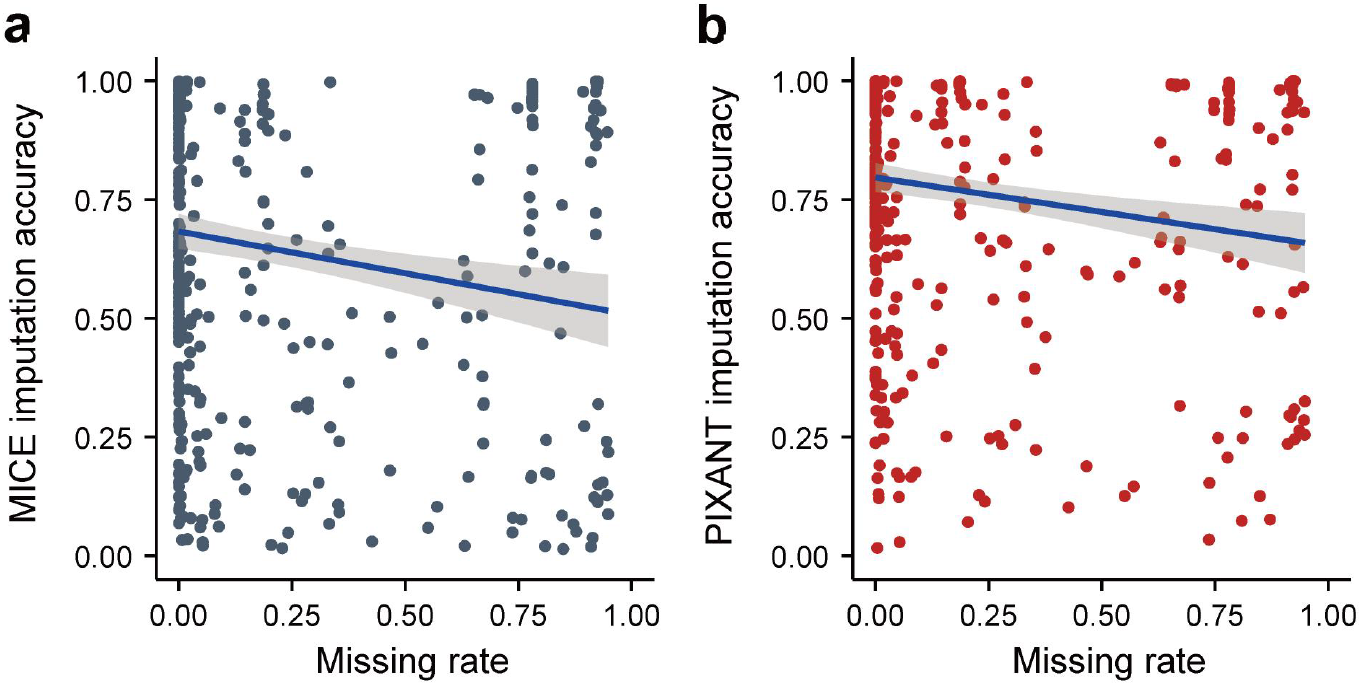
The impact of missing rate on the imputation accuracies of MICE (a) and PIXANT (b). The x-axis represents the missing rate of phenotypes, the y-axis represents imputation accuracies of PIXANT and MICE, and each point represents a phenotype. Although the linear fitting of missing rate and imputation accuracy was significant (*p*-values of 0.00029 and 0.00028 for PIXANT and MICE, respectively), the correlations are limited (the correlation between them is −0.176 and −0.175).

**Extended Data Fig. 3.**
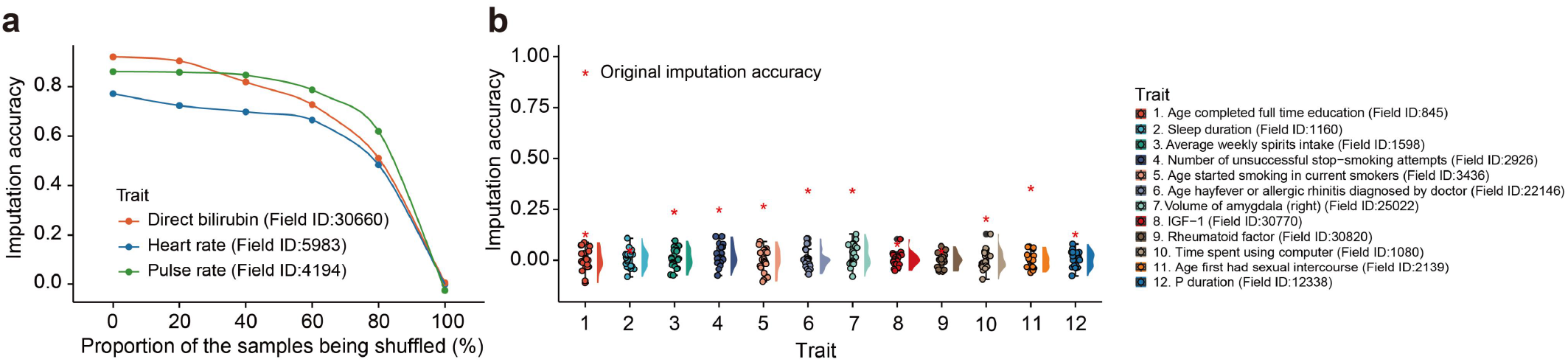
Permutation test as a negative control to assess imputation effects. (**a**) The impact of random shuffling reference phenotypes on imputation accuracy. The x-axis represents the proportion of reference phenotypes being shuffled, ranging from 0% to 100% in increments of 20%. Notably, for three traits, direct bilirubin (Field ID: 30660), heart rate (Field ID: 5983) and pulse rate (Field ID: 4194), the imputation accuracy gradually approaches to zero as the proportion of shuffled reference phenotypes increases to 100%, indicating that the accuracy is directly related to the non-random distribution of the reference phenotypes. (**b**) The empirical *p*-values for 12 selected phenotypes with imputation accuracies increases from 0.025 to 0.333. The empirical *p*-values were determined by 20 random permutations of the reference phenotypes. Higher accuracies (*r* =0.107, 0.333, etc.) correspond to *p*-values of zero, suggesting a significant imputation. In contrast, lower accuracies (*r* = 0.025, 0.028, 0.059) have *p*-values of 0.2, 0.4, and 0.05, respectively, not reaching the significance level.

**Extended Data Fig. 4.**
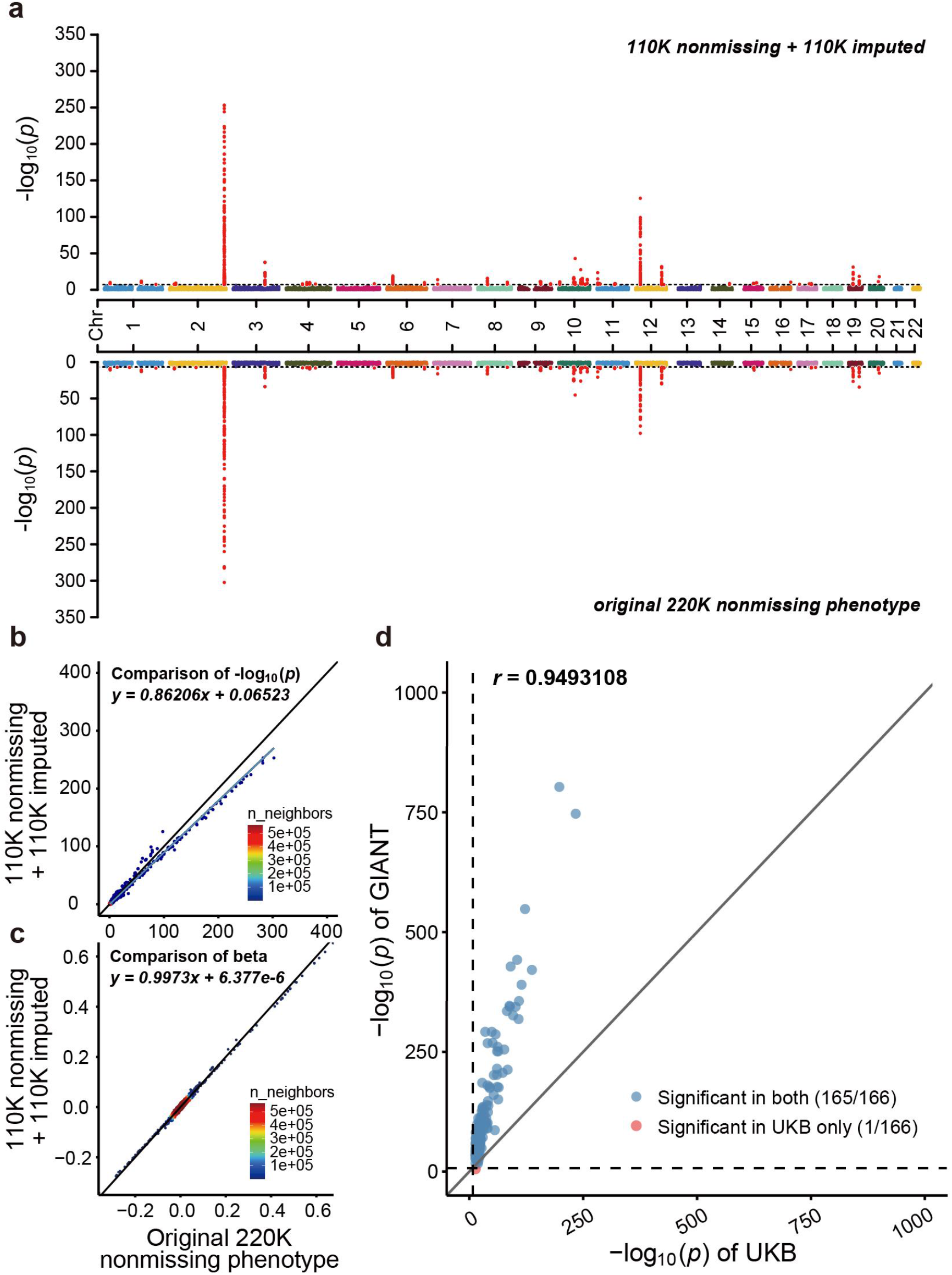
The replications in the real data analysis. We first assessed the repeatability of the GWAS discovered with bilirubin phenotype (Field ID 30660). We randomly masked 50% (110K) of known phenotypic records (220K in total) missing and imputed them with the remaining 110K know phenotype records using PIXANT, then compared the GWAS results of the above half-imputed dataset (110K nonmissing + 110K imputed phenotypes) with the complete dataset of 220K. (**a**) Visual comparison of the GWAS signals; (**b**) comparison of GWAS -log10 (*p*-values) for both data; and (**c**) comparison of GWAS beta values (regression coefficient) of association. While the hybrid dataset has marginally reduced power, as evidenced by a slight shift towards less significant *p*-values, the overall distribution remains consistent with that of using 220K full dataset. The findings suggest that PIXANT is effective in maintaining the generic GWAS findings even a substantial proportion of the data is imputed. (**d**) We further employed an independent dataset to validate the GWAS loci with imputed UKB data. We downloaded the human height summary statistics from GIANT consortium, which is generated with meta-analyses without UKB. We expected to replicate the 261 additional GWAS loci of human height with imputed UKB dataset. We extracted the lead SNPs of additional GWAS loci and compared the *p*-values with that of GIANT *p*-values. Among 261 lead SNPs, 166 has the corresponding *p*-value records in GIANT summary statistics file, and 165 are significant in GIANT while only one are not, where the *p*-value thresholds of GIANT (*p <* 3.63 ×10^-8^, determined with 0.05 divided by the number of SNPs) and PIXANT are represented with vertical and horizontal dotted lines, respectively; besides, the -log(*p*) values of both datasets are well correlated (*r* = 0.949), further demonstrating the validity of the identified GWAS loci of the imputed UKB dataset.

## References

1. Marx, V. Human phenotyping on a population scale. Nat. Methods 12, 711–714 (2015).

2. Schunkert, H. et al. Large-scale association analysis identifies 13 new susceptibility loci for coronary artery disease. Nat. Genet. 43, 333–338 (2011).

3. Voight, B. F. et al. Twelve type 2 diabetes susceptibility loci identified through large-scale association analysis. Nat. Genet. 42, 579–589 (2010).

4. Bycroft, C. et al. The UK Biobank resource with deep phenotyping and genomic data. Nature 562, 203–209 (2018).

5. Craig, J. E. et al. Multitrait analysis of glaucoma identifies new risk loci and enables polygenic prediction of disease susceptibility and progression. Nat. Genet. 52, 160–166 (2020).

6. Wray, N. R. et al. Genome-wide association analyses identify 44 risk variants and refine the genetic architecture of major depression. Nat. Genet. 50, 668–681 (2018).

7. Kemp, J. P. et al. Identification of 153 new loci associated with heel bone mineral density and functional involvement of GPC6 in osteoporosis. Nat. Genet. 49, 1468–1475 (2017).

8. Astle, W. J. et al. The Allelic Landscape of Human Blood Cell Trait Variation and Links to Common Complex Disease. Cel 167, 1415-1429.e19 (2016).

9. Dahl, A. et al. A multiple-phenotype imputation method for genetic studies. Nat. Genet. 48, 466–472 (2016).

10. Buuren, S. van & Groothuis-Oudshoorn, K. mice : Multivariate Imputation by Chained Equations in R. J. Stat. Softw. 45, 1–67 (2011).

11. Stekhoven, D. J. & Buhlmann, P. MissForest--non-parametric missing value imputation for mixed-type data. Bioinformatics 28, 112–118 (2012).

12. Schwarz, D. F., König, I. R. & Ziegler, A. On safari to Random Jungle: a fast implementation of Random Forests for high-dimensional data. Bioinformatics 26, 1752–1758 (2010).

13. Jin, H., Jung, S. & Won, S. missForest with feature selection using binary particle swarm optimization improves the imputation accuracy of continuous data. GenesGenomics 44, 651–658 (2022).

14. Zhou, X. & Stephens, M. Genome-wide efficient mixed-model analysis for association studies. Nat. Genet. 44, 821–824 (2012).

15. Zhou, X. & Stephens, M. Efficient multivariate linear mixed model algorithms for genome-wide association studies. Nat. Methods 11, 407–409 (2014).

16. Runcie, D. E. & Crawford, L. Fast and flexible linear mixed models for genome-wide genetics. PLOS Genet. 15, e1007978 (2019).

17. Knuth, D. E. TheArtofTheArtofComputerProgramming, Volume 2. (Addison-Wesley, Reading, 1985).

18. Wright, M. N. & Ziegler, A. ranger: A Fast Implementation of Random Forests for High Dimensional Data in C++ and R. J. Stat. Softw. 77, 1–17 (2017).

19. Yang, J. et al. Common SNPs explain a large proportion of the heritability for human height. Nat. Genet. 42, 565–569 (2010).

20. VanRaden, P. M. et al. Invited Review: Reliability of genomic predictions for North American Holstein bulls. J. DairySci. 92, 16–24 (2009).

21. Purcell, S. et al. PLINK: A Tool Set for Whole-Genome Association and Population-Based Linkage Analyses. Am. J. Hum. Genet. 81, 559–575 (2007).

22. Devlin, B., Roeder, K. & Wasserman, L. Genomic Control, a New Approach to Genetic-Based Association Studies. Theor. Popul. Biol. 60, 155–166 (2001).

23. Jordan, E. et al. Evidence-Based Assessment of Genes in Dilated Cardiomyopathy. Circulation 144, 7–19 (2021).

24. Mazzarotto, F. et al. Reevaluating the Genetic Contribution of Monogenic Dilated Cardiomyopathy. Circulation 141, 387–398 (2020).

25. Augusto, J. B. et al. Dilated cardiomyopathy and arrhythmogenic left ventricular cardiomyopathy: a comprehensive genotype-imaging phenotype study. Eur. HeartJ.-Cardiovasc. Imaging 21, 326–336 (2020).

26. Schroer, A. K. et al. Cadherin-11 blockade reduces inflammation-driven fibrotic remodeling and improves outcomes after myocardial infarction. JCIInsight 4, e131545 (2019).

27. Bowler, M. A. et al. Cadherin-11 as a regulator of valve myofibroblast mechanobiology. Am. J. lshoyioHr.-PeatCc. Phl. 315, H1614–H1626 (2018).

28. Ramírez, J. et al. Thirty loci identified for heart rate response to exercise and recovery implicate autonomic nervous system. Nat. Commun. 9, 1947 (2018).

29. Global BPgen Consortium et al. Identification of heart rate–associated loci and their effects on cardiac conduction and rhythm disorders. Nat. Genet. 45, 621–631 (2013).

30. Bezzina, C. R. et al. Common variants at SCN5A-SCN10A and HEY2 are associated with Brugada syndrome, a rare disease with high risk of sudden cardiac death. Nat. Genet. 45, 1044–1049 (2013).

31. van de Vegte, Y. J., Tegegne, B. S., Verweij, N., Snieder, H. & van der Harst, P. Genetics and the heart rate response to exercise. Cel. Mol. LifeSci. 76, 2391–2409 (2019).

32. Verstraeten, A., Luyckx, I. & Loeys, B. Aetiology and management of hereditary aortopathy. Nat. Rev. Cardiol. 14, 197–208 (2017).

33. Yang, J. et al. RGS6, a Modulator of Parasympathetic Activation in Heart. Circ. Res. 107, 1345–1349 (2010).

34. Wydeven, N., Posokhova, E., Xia, Z., Martemyanov, K. A. & Wickman, K. RGS6, but Not RGS4, Is the Dominant Regulator of G Protein Signaling (RGS) Modulator of the Parasympathetic Regulation of Mouse Heart Rate. J. Biol. Chem. 289, 2440–2449 (2014).

35. Ma, S.-L., Li, A.-J., Hu, Z.-Y.Shang, F.-S. & Wu, M.-C. Co-expression of the carbamoyl-phosphate synthase 1 gene and its long non-coding RNA correlates with poor prognosis of patients with intrahepatic cholangiocarcinoma. Mol. Med. Rep. 12, 7915–7926 (2015).

36. Abdel Ghany, E. A. G., Hussain, N. F. & Botros, S. K. A. Glutathione S-Transferase Gene Polymorphisms in Neonatal Hyperbilirubinemia. J. Investig. Med. 60, 18–22 (2012).

37. Muslu, N. et al. Are glutathione S-transferase gene polymorphisms linked to neonatal jaundice? Eur. J. Pediatr. 167, 57–61 (2007).

38. Yengo, L. et al. A saturated map of common genetic variants associated with human height. Nature 610, 704–712 (2022).

39. Hormozdiari, F. et al. Imputing Phenotypes for Genome-wide Association Studies. Am. J. Hum. Genet. 99, 89–103 (2016).

40. GTEx Consortium. Human genomics. The Genotype-Tissue Expression (GTEx) pilot analysis: multitissue gene regulation in humans. Science 348, 648–660 (2015).

41. Melé, M. et al. Human genomics. The human transcriptome across tissues and individuals. Science 348, 660–665 (2015).

42. Rivas, M. A. et al. Effect of predicted protein-truncating genetic variants on the human transcriptome. Science 348, 666–669 (2015).

43. Eraslan, G. et al. Single-nucleus cross-tissue molecular reference maps toward understanding disease gene function. Science 376, eabl4290 (2022).

44. Yin, X. et al. Integrating transcriptomics, metabolomics, and GWAS helps reveal molecular mechanisms for metabolite levels and disease risk. Am. J. Hum. Genet. 109, 1727–1741 (2022).

45. Park, P. J. ChIP–seq: advantages and challenges of a maturing technology. Nat. Rev. Genet. 10, 669–680 (2009).

46. Buenrostro, J. D., Giresi, P. G., Zaba, L. C., Chang, H. Y. & Greenleaf, W. J. Transposition of native chromatin for fast and sensitive epigenomic profiling of open chromatin, DNA-binding proteins and nucleosome position. Nat. Methods 10, 1213–1218 (2013).

47. Ziller, M. J., Hansen, K. D., Meissner, A. & Aryee, M. J. Coverage recommendations for methylation analysis by whole-genome bisulfite sequencing. Nat. Methods 12, 230–232 (2015).

48. Wang, Q. et al. Tagmentation-based whole-genome bisulfite sequencing. Nat. Protoc. 8, 2022–2032 (2013).

49. Wang, J. et al. Imputing Gene Expression in Uncollected Tissues Within and Beyond GTEx. Am. J. Hum. Genet. 98, 697–708 (2016).

50. Schafer, J. L. & Yucel, R. M. Computational Strategies for Multivariate Linear Mixed-Effects Models With Missing Values. J. Comput. Graph. Stat. 11, 437–457 (2002).

51. Yang, J. et al. Common SNPs explain a large proportion of the heritability for human height. Nat. Genet. 42, 565–569 (2010).

52. Privé, F., Luu, K., Blum, M. G. B., McGrath, J. J. & Vilhjálmsson, B. J. Effcient toolkit implementing best practices for principal component analysis of population genetic data. Bioinformatics 36, 4449–4457 (2020).

53. Privé, F., Aschard, H., Ziyatdinov, A. & Blum, M. G. B. Efficient analysis of large-scale genome-wide data with two R packages: bigstatsr and bigsnpr. Bioinformatics 34, 2781–2787 (2018).

54. Li, Y. et al. Genomic analyses of an extensive collection of wild and cultivated accessions provide new insights into peach breeding history. GenomeBiol. 20, 1–18 (2019).

