## Supplemental Tables for "Rapid and accurate multi-phenotype imputation for millions of individuals"

Supplementary Material

Lin-Lin Gu[1], Hong-Shan Wu[1], Tian-Yi Liu[1], Yong-Jie Zhang[1], Jing-Cheng He[2], Xiao-Lei Liu[3], Zhi-Yong Wang[1], Guo-Bo Chen[4,5*], Dan Jiang[1*] and Ming Fang[1*]

[1] Key Laboratory of Healthy Mariculture for the East China Sea, Ministry of Agriculture and Rural Affairs & Fisheries college, Jimei University, Xiamen, Fujian, People's Republic of China

[2] Center for Data Science, School of Mathematical Sciences, Zhejiang University, Hangzhou, Zhejiang, People's Republic of China

[3] Key Laboratory of Agricultural Animal Genetics, Breeding and Reproduction, Ministry of Education & College of Animal Science and Technology, Huazhong Agricultural University, Wuhan, Hubei, People's Republic of China

[4] Center for General Practice Medicine, Department of General Practice Medicine, Clinical Research Institute, Zhejiang Provincial People's Hospital, People's Hospital of Hangzhou Medical College, Hangzhou, Zhejiang, People's Republic of China

[5] Key Laboratory of Endocrine Gland Diseases of Zhejiang Province, Hangzhou, Zhejiang, People's Republic of China

***Correspondence:**

Ming Fang<>;

Dan Jiang<>;

Guo-Bo Chen<>

**Supplementary Tab. 1** Brief summary of the methods applied to simulated and real data sets.

| Method | Description and properties | Reference |
|---|---|---|
| PIXANT | A phenotype imputation with mixed fast random forest. | This paper |
| PHENIX | Bayesian multivariate mixed model fitted via variational Bayes. | [1] |
| missForest | non-parametric missing value imputation for mixed-type data. | [2] |
| MICE | Multivariate imputation by chained equations. | [3] |
| LMM | Single-trait linear mixed model with estimated BLUP used to impute missing values; ignores covariance between phenotypes. | [4–6] |

**Supplementary Tab. 2** The imputation accuracies of different methods with different sample size.

| Method | Sample size | | | | | |
|---|---|---|---|---|---|---|
| | 100 | 300 | 500 | 700 | 900 | 1100 |
| | Imputation accuracy | Imputation accuracy | Imputation accuracy | Imputation accuracy | Imputation accuracy | Imputation accuracy |
| PIXANT | 0.3018±0.0059 | 0.5884±0.0030 | 0.6340±0.0020 | 0.6511±0.0018 | 0.6579±0.0016 | 0.6657±0.0015 |
| PHENIX | 0.3976±0.0057 | 0.5760±0.0025 | 0.6178±0.0020 | 0.6367±0.0019 | 0.6445±0.0016 | 0.6526±0.0015 |
| missForest | 0.2876±0.0060 | 0.3930±0.0029 | 0.4356±0.0026 | 0.4567±0.0022 | 0.4696±0.0015 | 0.4814±0.0016 |
| MICE | 0.2595±0.0053 | 0.4349±0.0032 | 0.4554±0.0026 | 0.4679±0.0024 | 0.4684±0.0023 | 0.4705±0.0021 |
| LMM | 0.0847±0.0054 | 0.0932±0.0036 | 0.0966±0.0033 | 0.1000±0.0026 | 0.1052±0.0023 | 0.1045±0.0022 |

**Note:** ± Represent the standard deviation (SD).

**Supplementary Tab. 3** The imputation accuracies of different methods with different heritability.

| Method | Heritability | | | | | |
|---|---|---|---|---|---|---|
| | 0.1 | 0.2 | 0.4 | 0.6 | 0.8 | 1.0 |
| | Imputation accuracy | Imputation accuracy | Imputation accuracy | Imputation accuracy | Imputation accuracy | Imputation accuracy |
| PIXANT | 0.7785±0.0012 | 0.7050±0.0017 | 0.6334±0.0021 | 0.6331±0.0021 | 0.7043±0.0017 | 0.9160±0.0008 |
| PHENIX | 0.7665±0.0012 | 0.6925±0.0016 | 0.6185±0.0019 | 0.6177±0.0019 | 0.6902±0.0016 | 0.9112±0.0007 |
| missForest | 0.5549±0.0018 | 0.4999±0.0020 | 0.4353±0.0021 | 0.4340±0.0020 | 0.4963±0.0020 | 0.6091±0.0017 |
| MICE | 0.6479±0.0017 | 0.5487±0.0022 | 0.4557±0.0025 | 0.4596±0.0025 | 0.5436±0.0021 | 0.9116±0.0011 |
| LMM | 0.0280±0.0023 | 0.0476±0.0023 | 0.0757±0.0026 | 0.0971±0.0028 | 0.1146±0.0030 | 0.1284±0.0031 |

**Note:** ± Represent the standard deviation (SD).

**Supplementary Tab. 4** The imputation accuracies of different methods with different number of phenotypes.

| Method | Phenotype number | | | | | |
|---|---|---|---|---|---|---|
| | 20 | 40 | 60 | 80 | 100 | 120 |
| | Imputation accuracy | Imputation accuracy | Imputation accuracy | Imputation accuracy | Imputation accuracy | Imputation accuracy |
| PIXANT | 0.2251±0.0048 | 0.3286±0.0028 | 0.4053±0.0023 | 0.4686±0.0018 | 0.5233±0.0017 | 0.5726±0.0015 |
| PHENIX | 0.2375±0.0047 | 0.3275±0.0029 | 0.3951±0.0024 | 0.4555±0.0019 | 0.5065±0.0017 | 0.5554±0.0014 |
| missForest | 0.1854±0.0048 | 0.2527±0.0032 | 0.2874±0.0025 | 0.3134±0.0021 | 0.3320±0.0018 | 0.3474±0.0018 |
| MICE | 0.0847±0.0043 | 0.1585±0.0033 | 0.2310±0.0024 | 0.2997±0.0021 | 0.3652±0.0021 | 0.4299±0.0017 |
| LMM | 0.1002±0.0044 | 0.0987±0.0034 | 0.0975±0.0026 | 0.0967±0.0024 | 0.0970±0.0022 | 0.0973±0.0020 |

**Note:** ± Represent the standard deviation (SD).

**Supplementary Tab. 5** The imputation accuracies of different methods with different sample relatedness.

| **Method** | **Sample relatedness** | | | | | |
|---|---|---|---|---|---|---|
| | **0.02** | **0.2** | **0.4** | **0.6** | **0.8** | **0.9** |
| | **Imputation accuracy** | **Imputation accuracy** | **Imputation accuracy** | **Imputation accuracy** | **Imputation accuracy** | **Imputation accuracy** |
| PIXANT | 0.6331±0.0021 | 0.6731±0.0021 | 0.7212±0.0024 | 0.7749±0.0024 | 0.8413±0.0021 | 0.8862±0.0017 |
| PHENIX | 0.6177±0.0019 | 0.6620±0.0020 | 0.7134±0.0024 | 0.7701±0.0025 | 0.8386±0.0022 | 0.8838±0.0017 |
| missForest | 0.4338±0.0020 | 0.5043±0.0025 | 0.5881±0.0034 | 0.6700±0.0036 | 0.7512±0.0033 | 0.7932±0.0030 |
| MICE | 0.4596±0.0025 | 0.5057±0.0027 | 0.5681±0.0032 | 0.6426±0.0034 | 0.7395±0.0032 | 0.8108±0.0026 |
| LMM | 0.0971±0.0028 | 0.3354±0.0042 | 0.4769±0.0048 | 0.5859±0.0049 | 0.6787±0.0046 | 0.7211±0.0043 |

**Note:** ± Represent the standard deviation (SD).

**Supplementary Tab. 6** The comparison of runtime (in hours) and memory usage (MB) of PHENIX and PIXANT for different sample size.

| **Sample size** | **Runtime (h)** | | **Memory usage (MB)** | |
|---|---|---|---|---|
| | **PHENIX** | **PIXANT** | **PHENIX** | **PIXANT** |
| 1000 | 0.06 | 0.04 | 670.49 | 1.71 |
| 5000 | 0.60 | 0.08 | 1942.51 | 2.23 |
| 10,000 | 2.02 | 0.15 | 3627.67 | 2.60 |
| 15,000 | 4.44 | 0.23 | 5435.97 | 3.87 |
| 20,000 | 7.38 | 0.29 | 8052.91 | 5.12 |

**Note:** Computing performance tests were performed in a Red Hat Enterprise Linux sever with 2.20 GHz Intel(R) Xeon(R) Silver 4114 CPU, and 125 GB memory.

**Supplementary Tab. 7** The comparison of runtime (in hours) and memory usage (MB) of PHENIX and PIXANT for different number of phenotypes.

| Phenotype number | Runtime (h) | | Memory usage (Mb) | |
|---|---|---|---|---|
| | PHENIX | PIXANT | PHENIX | PIXANT |
| 10 | 1.89 | 0.08 | 815.53 | 2.60 |
| 15 | 1.90 | 0.11 | 889.99 | 2.90 |
| 20 | 1.91 | 0.13 | 1406.57 | 3.24 |
| 25 | 1.95 | 0.14 | 2926.26 | 3.44 |
| 30 | 2.02 | 0.15 | 3627.68 | 3.88 |

**Note:** Computing performance tests were performed in a Red Hat Enterprise Linux sever with 2.20 GHz Intel(R) Xeon(R) Silver 4114 CPU, and 125 Gb memory.

**Supplementary Tab. 8** Genome wide Significant Associations of extra Single Nucleotide Polymorphisms (SNPs) with Heart rate (**Field ID: 5983**) and Direct bilirubin (**Field ID: 30660**).

| Trait | GWAS site | Lead SNP | Position | *p*-value (unimputed/imputed) | Gene (Distance, bp) |
|---|---|---|---|---|---|
| Heart rate | chr1:44505996-45505996 | rs272570 | 45005996 | 0.00204664/3.86E-10 | ***RNF220***[7–9] (0) |
| Heart rate | chr11:61057803-62057803 | rs102275 | 61557803 | 3.11E-05/1.07E-13 | ***FADS2***[8] (2649), ***FADS1***[10,11] (9296) |
| Heart rate | chr5:122181834-123181834 | rs1047440 | 122681834 | 0.000966971/7.56E-09 | ***CEP120***[12] (0) |
| Heart rate | chr12:33033374-34033374 | rs10732554 | 33533374 | 2.05E-06/1.13E-24 | ***SYT10***[7–10,13] (0) |
| Heart rate | chr18:25266218-26266218 | rs11083258 | 25766218 | 0.004168892/5.96E-10 | ***CDH2***[8] (8808) |
| Heart rate | chr3:49038799-50038799 | rs11130199 | 49538799 | 0.000117932/1.42E-08 | ***DAG1***[8] (0) |
| Heart rate | chr12:37974090-38974090 | rs11182193 | 38474090 | 0.002751144/4.24E-08 | ***ALG10B***[7] (236290) |
| Heart rate | chr15:73162855-74162855 | rs11630367 | 73662855 | 3.39E-06/2.30E-22 | ***HCN4***[8,10,13,14] (1250), ***NEO1***[8,13] (65308) |
| Heart rate | chr6:133714525-134714525 | rs12190287 | 134214525 | 0.000251269/1.83E-08 | ***TCF21***[15–17] (0) |
| Heart rate | chr12:19968094-20968094 | rs12581906 | 20468094 | 0.002775923/1.47E-08 | ***PDE3A***[8] (54085) |

| | | | | | |
|---|---|---|---|---|---|
| Heart rate | chr2:27230940-28230940 | rs1260326 | 27730940 | 0.000508897/3.25E-10 | ***GCKR***[8] (0) |
| Heart rate | chr1:207524820-208524820 | rs12731740 | 208024820 | 1.83E-05/1.60E-20 | ***CD46***[8,10] (55962), ***CD34***[8,11] (32774), ***PLXNA2***[18](170767) |
| Heart rate | chr7:100006381-101006381 | rs13226502 | 100506381 | 3.38E-06/9.08E-34 | ***ACHE***[7–10] (11787), ***UFSP1***[8,9] (19042), ***TRIP6***[9] (35305) |
| Heart rate | chr2:219799018-220799018 | rs13386459 | 220299018 | 0.004147206/1.58E-09 | ***SPEG***[8] (550) |
| Heart rate | chr14:72385471-73385471 | rs17180489 | 72885471 | 7.14E-05/8.28E-20 | ***RGS6***[7–9,13,19] (0) |
| Heart rate | chr1:216231498-217231498 | rs17668665 | 216731498 | 0.022556358/4.57E-08 | ***ESRRG***[18,20,21] (0) |
| Heart rate | chr7:93050447-94050447 | rs180238 | 93550447 | 0.000687127/5.14E-15 | ***GNG11***[8–10,13] (564) |
| Heart rate | chr7:115933527-116933527 | rs193688 | 116433527 | 0.008190843/3.83E-08 | ***CAV2***[7,9] (284932) |
| Heart rate | chr5:136919989-137919989 | rs2040862 | 137419989 | 0.000423035/1.61E-14 | ***WNT8A***[22] (0), ***GFRA3***[23] (168079) |
| Heart rate | chr12:1678661- 2678661 | rs2238018 | 2178661 | 0.000261796/2.92E-13 | ***CACNA1C***[8] (0) |
| Heart rate | chr2:231763127-232763127 | rs2290130 | 232263127 | 3.05E-05/6.21E-12 | ***B3GNT7***[8,10] (0) |
| Heart rate | chr3:49674197-50674197 | rs2624847 | 50174197 | 0.005074611/5.17E-10 | ***GNAI2***[24] (89527) |
| Heart rate | chr16:15418096-16418096 | rs2733855 | 15918096 | 7.38E-05/1.68E-09 | ***MYH11***[8] (0) |
| Heart rate | chr11:128268938-129268938 | rs2846700 | 128768938 | 3.52E-05/6.03E-11 | ***KCNJ5***[8] (0) |
| Heart rate | chr14:85292358-86292358 | rs35195423 | 85792358 | 2.14E-07/1.81E-19 | ***FLRT2***[8,10] (204130) |
| Heart rate | chr6:7043468-8043468 | rs3823183 | 7543468 | 0.04158108/7.49E-08 | ***DSP***[25–27] (0) |
| Heart rate | chr3:38267315-39267315 | rs6801957 | 38767315 | 0.000925654/1.52E-08 | ***SCN10A***[7,28–30] (0) |
| Heart rate | chr4:148474602-149474602 | rs6845865 | 148974602 | 0.001566177/1.73E-11 | ***NR3C2***[31,32] (25311) |
| Heart rate | chr7:136102131-137102131 | rs73158724 | 136602131 | 0.004630562/7.10E-08 | ***CHRM2***[7–10] (0) |
| Heart rate | chr18:33706449-34706449 | rs79026601 | 34206449 | 0.011565212/1.55E-08 | ***FHOD3***[8] (0) |
| Heart rate | chr16:64779257-65779257 | rs9921437 | 65279257 | 0.000424767/2.31E-11 | ***CDH11***[8] (119242) |
| Direct bilirubin | chr2:211040507-212040507 | rs1047891 | 211540507 | 6.09E-05/1.90E-09 | ***CPS1***[33](0) |
| Direct bilirubin | chr1:109732983-110732983 | rs140584594 | 110232983 | 1.43E-06/5.57E-10 | ***GSTM1***[34,35](0) |
| Direct bilirubin | chr1:55005647-56005647 | rs11591147 | 55505647 | 6.84E-06/3.18E-10 | ***PCSK9***[36–38](0) |
| Direct bilirubin | chr3:170225542-171225542 | rs10513686 | 170725542 | 8.05E-06/1.67E-09 | ***SLC2A2***[39](0) |

| | | | | | |
|---|---|---|---|---|---|
| Direct bilirubin | chr4:144420596-145420596 | rs7683365 | 144920596 | 1.10E-05/7.08E-08 | ***GYPB***[40](0) |
| Direct bilirubin | chr6:34872524-35872524 | rs2267667 | 35372524 | 1.83E-05/3.80E-08 | ***PPARD***[41](0) |
| Direct bilirubin | chr10:113413222-114413222 | rs2297991 | 113913222 | 1.19E-07/5.36E-08 | ***GPAM***[38,42](0) |
| Direct bilirubin | chr15:43320717-44320717 | rs55707100 | 43820717 | 6.45E-06/6.16E-10 | ***MAP1A***[43](0) |
| Direct bilirubin | chr22:36969591-37969591 | rs4820268 | 37469591 | 8.75E-07/2.48E-15 | ***TMPRSS6***[44,45](0) |

**Note:** A genome-wide significance threshold (α = 0.05 / 563,675; ***p*-value** $< \mathbf{8.87 \times 10^{-8}}$).

### Supplementary reference


1. Dahl, A. *et al.* A multiple-phenotype imputation method for genetic studies. *Nat. Genet.* **48**, 466–472 (2016).
2. Stekhoven, D. J. & Buhlmann, P. MissForest--non-parametric missing value imputation for mixed-type data. *Bioinformatics* **28**, 112–118 (2012).
3. Buuren, S. van & Groothuis-Oudshoorn, K. mice : Multivariate Imputation by Chained Equations in R. *J. Stat. Softw.* **45**, 1–67 (2011).
4. Zhou, X. & Stephens, M. Genome-wide efficient mixed-model analysis for association studies. *Nat. Genet.* **44**, 821–824 (2012).
5. Zhou, X. & Stephens, M. Efficient multivariate linear mixed model algorithms for genome-wide association studies. *Nat. Methods* **11**, 407–409 (2014).
6. Runcie, D. E. & Crawford, L. Fast and flexible linear mixed models for genome-wide genetics. *PLOS Genet.* **15**, e1007978 (2019).
7. Ramírez, J. *et al.* Thirty loci identified for heart rate response to exercise and recovery implicate autonomic nervous system. *Nat. Commun.* **9**, 1947 (2018).
8. Eppinga, R. N. *et al.* Identification of genomic loci associated with resting heart rate and shared genetic predictors with all-cause mortality. *Nat. Genet.* **48**, 1557–1563 (2016).
9. Verweij, N., van de Vegte, Y. J. & van der Harst, P. Genetic study links components of the autonomous nervous system to heart-rate profile during exercise. *Nat. Commun.* **9**, 898 (2018).
10. den Hoed, M. *et al.* Identification of heart rate-associated loci and their effects on cardiac conduction and rhythm disorders. *Nat. Genet.* **45**, 621–631 (2013).

11. Eijgelsheim, M. *et al.* Genome-wide association analysis identifies multiple loci related to resting heart rate. *Hum. Mol. Genet.* **19**, 3885–3894 (2010).

12. Vasan, R. S. *et al.* Genetic variants associated with cardiac structure and function: a meta-analysis and replication of genome-wide association data. *JAMA* **302**, 168–178 (2009).

13. Nolte, I. M. *et al.* Genetic loci associated with heart rate variability and their effects on cardiac disease risk. *Nat. Commun.* **8**, 15805 (2017).

14. Baruscotti, M. *et al.* A gain-of-function mutation in the cardiac pacemaker HCN4 channel increasing cAMP sensitivity is associated with familial Inappropriate Sinus Tachycardia. *Eur. Heart J.* **38**, 280–288 (2017).

15. Zhao, Q. *et al.* Molecular mechanisms of coronary disease revealed using quantitative trait loci for TCF21 binding, chromatin accessibility, and chromosomal looping. *Genome Biol.* **21**, 135 (2020).

16. Lu, X. *et al.* Genome-wide association study in Han Chinese identifies four new susceptibility loci for coronary artery disease. *Nat. Genet.* **44**, 890–894 (2012).

17. Schunkert, H. *et al.* Large-scale association analysis identifies 13 new susceptibility loci for coronary artery disease. *Nat. Genet.* **43**, 333–338 (2011).

18. Parsa, A. *et al.* Hypertrophy-associated polymorphisms ascertained in a founder cohort applied to heart failure risk and mortality. *Clin. Transl. Sci.* **4**, 17–23 (2011).

19. Posokhova, E., Wydeven, N., Allen, K. L., Wickman, K. & Martemyanov, K. A. RGS6/Gβ5 complex accelerates IKACh gating kinetics in atrial myocytes and modulates parasympathetic regulation of heart rate. *Circ. Res.* **107**, 1350–1354 (2010).

20. Simino, J., Sung, Y. J., Kume, R., Schwander, K. & Rao, D. C. Gene-alcohol interactions identify several novel blood pressure loci including a promising locus near SLC16A9. *Front. Genet.* **4**, 277 (2013).

21. Sakamoto, T. *et al.* The nuclear receptor ERR cooperates with the cardiogenic factor GATA4 to orchestrate cardiomyocyte maturation. *Nat. Commun.* **13**, 1991 (2022).

22. Ellinor, P. T. *et al.* Meta-analysis identifies six new susceptibility loci for atrial fibrillation. *Nat. Genet.* **44**, 670–675 (2012).

23. Arking, D. E. *et al.* Genetic association study of QT interval highlights role for calcium signaling pathways in myocardial repolarization. *Nat. Genet.* **46**, 826–836 (2014).

24. Lerman, B. B. *et al.* Right ventricular outflow tract tachycardia due to a somatic cell mutation in G protein subunitalphai2. *J. Clin. Invest.* **101**, 2862–2868 (1998).

25. Bariani, R. *et al.* Clinical profile and long-term follow-up of a cohort of patients with desmoplakin cardiomyopathy. *Heart Rhythm* **19**, 1315–1324 (2022).

26. Karvonen, V. *et al.* A novel desmoplakin mutation causes dilated cardiomyopathy with palmoplantar keratoderma as an early clinical sign. *J. Eur. Acad. Dermatol. Venereol. JEADV* **36**, 1349–1358 (2022).

27. Hylind, R. *et al.* Cardiocutaneous Features of Autosomal Dominant Desmoplakin-Associated Arrhythmogenic Cardiomyopathy. *Circ. Genomic Precis. Med.* **13**, e003081 (2020).

28. Sano, M. *et al.* Genome-wide association study of electrocardiographic parameters identifies a new association for PR interval and confirms previously reported associations. *Hum. Mol. Genet.* **23**, 6668–6676 (2014).

29. Holm, H. *et al.* Several common variants modulate heart rate, PR interval and QRS duration. *Nat. Genet.* **42**, 117–122 (2010).

30. Chambers, J. C. *et al.* Genetic variation in SCN10A influences cardiac conduction. *Nat. Genet.* **42**, 149–152 (2010).

31. Rossier, M. F. The Cardiac Mineralocorticoid Receptor (MR): A Therapeutic Target Against Ventricular Arrhythmias. *Front. Endocrinol.* **12**, 694758 (2021).

32. Parker, B. M. *et al.* Novel Insights into the Crosstalk between Mineralocorticoid Receptor and G Protein-Coupled Receptors in Heart Adverse Remodeling and Disease. *Int. J. Mol. Sci.* **19**, 3764 (2018).

33. Ma, S.-L., Li, A.-J., Hu, Z.-Y., Shang, F.-S. & Wu, M.-C. Co-expression of the carbamoyl-phosphate synthase 1 gene and its long non-coding RNA correlates with poor prognosis of patients with intrahepatic cholangiocarcinoma. *Mol. Med. Rep.* **12**, 7915–7926 (2015).

34. Abdel Ghany, E. A. G., Hussain, N. F. & Botros, S. K. A. Glutathione S-Transferase Gene Polymorphisms in Neonatal Hyperbilirubinemia. *J. Investig. Med.* **60**, 18–22 (2012).

35. Muslu, N. *et al.* Are glutathione S-transferase gene polymorphisms linked to neonatal jaundice? *Eur. J. Pediatr.* **167**, 57–61 (2007).

36. Cohen, J. *et al.* Low LDL cholesterol in individuals of African descent resulting from frequent nonsense mutations in PCSK9. *Nat. Genet.* **37**, 161–165 (2005).

37. Abifadel, M. *et al.* Mutations in PCSK9 cause autosomal dominant hypercholesterolemia. *Nat. Genet.* **34**, 154–156 (2003).

38. Goga, A. & Stoffel, M. Therapeutic RNA-silencing oligonucleotides in metabolic diseases. *Nat. Rev. Drug Discov.* **21**, 417–439 (2022).

39. Morioka, S. *et al.* Efferocytosis induces a novel SLC program to promote glucose uptake and lactate release. *Nature* **563**, 714–718 (2018).

40. Saleh, R. M., Zefarina, Z., Che Mat, N. F., Chambers, G. K. & Edinur, H. A. Transfusion Medicine and Molecular Genetic Methods. *Int. J. Prev. Med.* **9**, 45 (2018).

41. Wickramasinghe, N. M. *et al.* PPARdelta activation induces metabolic and contractile maturation of human pluripotent stem cell-derived cardiomyocytes. *Cell Stem Cell* **29**, 559-576.e7 (2022).

42. Ng, S. W. K. *et al.* Convergent somatic mutations in metabolism genes in chronic liver disease. *Nature* **598**, 473–478 (2021).

43. Halpain, S. & Dehmelt, L. The MAP1 family of microtubule-associated proteins. *Genome Biol.* **7**, 224 (2006).

44. An, P. *et al.* TMPRSS6, but not TF, TFR2 or BMP2 variants are associated with increased risk of iron-deficiency anemia. *Hum. Mol. Genet.* **21**, 2124–2131 (2012).

45. Gan, W. *et al.* Association of TMPRSS6 polymorphisms with ferritin, hemoglobin, and type 2 diabetes risk in a Chinese Han population. *Am. J. Clin. Nutr.* **95**, 626–632 (2012).
